# Quorum sensing in *Bacillus subtilis* slows down biofilm formation by enabling sporulation bet hedging

**DOI:** 10.1101/768671

**Authors:** Mihael Spacapan, Tjaša Danevčič, Polonca Štefanic, Ines Mandic-Mulec

**Author notes:** Address correspondence to Ines Mandic-Mulec,.

## Abstract

1.2

The ComQXPA quorum sensing (QS) system of *Bacillus subtilis*, a Gram-positive, industrially relevant, endospore forming bacterium, promotes surfactin production. This lipopeptide increases transcription of several genes involved in biofilm matrix synthesis via the Spo0A-P master regulator. We hypothesized that the inactivation of the QS system will therefore result in decreased rates of floating biofilm formation. We find that this is not the case and that the QS deficient mutant forms pellicles with a faster rate and produces more biofilm matrix components than the wild type. As Spo0A-P is the master regulator of sporulation initiation we hypothesized that the ComQXPA dependent signaling promotes sporulation and consequently slows the growth rate of the wild type strain. Indeed, our results confirm that cells with the inactive QS initiate endospore formation in biofilms later and more synchronously than the wild type, as evidenced by spore frequencies and the P_*spoIIQ*_ promoter activity. We argue, that the QS system acts as a switch that promotes stochastic sporulation initiation and consequently bet hedging behavior. By committing a subpopulation of cells to sporulation early during growth, wild type population grows slower and produces thinner biofilms but also assures better survival under stressful conditions.

**IMPORTANCE:** *Bacillus subtilis* is widely employed model organism to study biofilm formation and sporulation in Gram-positive bacteria. The ComQXPA quorum sensing (QS) system indirectly increases the transcription of genes involved in biofilm matrix formation, which predicts a positive role of this QS in biofilm development Here we show that QS mutants actually form more matrix components per pellicle than the wild type and that their pellicles are thicker and form with a faster rate. We explain this, by showing that cells with an inactive QS exhibit a delay in sporulation entry, which is also more synchronous relative to the wild type. We argue, that the ComQXPA QS system acts as a switch that contributes to the stochastic sporulation initiation and though this path promotes bet hedging behavior. This finding is important in terms of “quorum quenching” strategies aiming to down modulate biofilm development through inhibition of QS signaling and underscores the richness of QS regulated phenotypic outcomes among bacterial species.

## 1.3 INTRODUCTION

Biofilms are multicellular groups encased in extracellular matrix that are believed to be the default mode of microbial growth in nature (1–3). Biofilm formation is often regulated by quorum sensing (QS) (4), a wide spread microbial communication system that coordinates bacterial gene expression in accordance with cell density (5). *Bacillus subtilis* is the most studied species in the genus *Bacillus* (6, 7), which has served over the years as an excellent model to investigate development of metabolically dormant and heat resistant spores (8), a model for biofilm development (9–12), division of labor (13–16) and for a variety of intra and interspecies social interactions (17). This Gram-positive spore former relies on peptide based QS system, encoded by the *comQXPA* gene cluster (18, 19), which is wide spread among Firmicutes (20). It controls transcriptional activity of several adaptive processes including synthesis of a lipopetide antibiotic surfactin (21–23), exoprotease production (24, 25), and competence development (26). The extracellular signaling peptide ComX is synthesized as a pre-peptide, cleaved and post-translationally modified by the isoprenoid transferase ComQ (21, 27). Therefore, the gene *comQ* is essential for a functional and active QS system. Mature and biologically active ComX accumulates extracellularly, where it interacts with the sensory histidine kinase ComP, which then phosphorylates ComA (28). ComA-P in its active phosphorylated form regulates transcription of several target genes (29–31). ComA-P also increases the transcription rates of the pleiotropic regulatory gene *degQ* (25, 30, 32), which ultimately increases the DegU phosphorylation rate (25). DegU-P is required for the activation of extracellular degradative enzyme production. This regulatory network is relevant also during biofilm development (33), where operational ComX dependent QS is required for exoprotease production (24).

In biofilms, cells are encased in a structure formed out of extracellular polysaccharides (Eps), proteins, and extracellular DNA (3); but the ratios of each biofilm constituent differ depending on the specific strain, media and growth conditions (34). In *B. subtilis* the *epsA-O* operon is involved in the production of the major polysaccharide component of the biofilm matrix (35), which is essential for development of floating biofilm (pellicle) (36). TasA, the major matrix protein, encoded by the *tapA-tasA-sipW* operon (hereafter referred to as the *tapA* operon), gives structural support to floating biofilm. Although TasA and TapA proteins are not essential for the formation of floating biofilms, the *tasA* mutant forms a less prominent structures (37, 38). The molecular regulation of the operons involved in the synthesis of the biofilm matrix components is very complex (39). Briefly, biofilm development is dependent on phosphorylation state of the major transcription factor, Spo0A (36), which is controlled by the phosphorelay of multiple histidine kinases (40). Spo0A-P activates transcription of the *sinI* operon (41, 42). SinI inihibts SinR (43) which acts as the main transcriptional repressor of the *epsA* and *tapA* operons (36, 44). All in all an intermediate amount of phosphorylated Spo0A is believed to increase the transcription of the biofilm matrix production operons, at very high Spo0A-P levels, however, cells start the spore formation process (44) during which they become less metabolically active. The second regulator affecting biofilm formation is DegU, gradual phosphorylation of this regulator is essential for floating biofilm formation (45), which in its phosphorylated form increases *bslA* transcription (46) and poly-γ-glutamate (*pgs*) operon transcription (47). Poly-γ-glutamate only plays a role during the formation of surface adhered biofilms (47). It has been also shown that DegU indirectly affects the phosphorylation state of Spo0A, shortening the time window of intermediate Spo0A phosphorylation, required to trigger synthesis of the extracellular matrix (48).

The ComQXPA QS system can affect the transcription of the biofilm matrix operons by increasing surfactin synthesis. Surfactin, by initiating potassium ion leakage, triggers the activity of KinC (49). This histidine kinase then through phosphorelay system increases the phosphorylation state of Spo0A (50). Thus the ComQXPA QS system indirectly increases the phosphorylation of Spo0A via the lipopeptide surfactin. QS and/or surfactin are therefore believed to promote biofilm formation (4, 12, 13, 16, 39, 51–53).

With this point in mind we present our hypotheses, which were tested by comparing different characteristics of floating biofilms formed by strains exhibiting either a wild type phenotype (wt) or a QS inactivated phenotype (Δ*comQ*), where this mutant fails to produce an active signaling peptide:

### Hypothesis 1

The inactivation of the QS system will lead to decreased surfactin production, and stunted floating biofilm development.

This hypothesis might appear self-evident, especially in the light of all the *a priori* knowledge in regards to the transcriptional regulation of the two major operons involved in biofilm formation. While surfactin production was indeed decreased in the Δ*comQ* floating biofilm, we observed that the floating biofilm formation was actually promoted. We therefore concluded that our data does not support hypothesis 1. We then tested two more additional hypotheses in an attempt to explain this unexpected phenomenon.

### Hypothesis 2

The ComQXPA QS system negatively regulates transcription of matrix producing genes (*epsA-O, tapA*) in the PS-216 *B. subtilis* strain at the single cell level.

Our results, however, did not support this hypothesis when transcription of these operons was quantified at the single cell level, which was in agreement with previously published results on QS regulated expression of *epsA-O* in the more widely investigated *B. subtilis* NCIB 3610 strain (49). In contrast, our results indicated higher cumulative production of matrix components when these were quantified per pellicle, suggesting that biofilms may harbor a higher number of metabolically active cells. To investigate this further we proposed and tested a third hypothesis.

### Hypothesis 3

A developing wild type floating biofilm contains more cells committed to sporulation during biofilm development compared to the Δ*comQ*

If the third hypothesis is true, then there are more cells producing biofilm matrix components at any given time point the Δ*comQ* floating biofilm relative to the wild type. Therefore, the difference in rates of biofilm formation between the mutant and the wild type may arise due to the absence of stochastic sporulation initiation. We found this hypothesis plausible, because Spo0A phosphorylation is also essential for the regulation of sporulation initiation, and because the QS inactivated strain had a faster growth rate than the wild type strain. All of our results were in accord with our third and final hypothesis.

## 1.4 RESULTS

### 1.4.1 Hypothesis 1

The inactivation of the QS system will lead to decreased surfactin production, and stunted floating biofilm development.

To test this hypothesis we first measured the fluorescence of entire floating biofilms (pellicles) formed by a strain exhibiting the wild type phenotype (wt) and a strain exhibiting a QS mutant phenotype (Δ*comQ*) carrying a *srfA* transcriptional reporter. In such reporter strains, the transcription of a fluorescent protein gene is controlled by a specific promoter of interest (In this case the *srfA* promoter, regulating the transcription of surfactin synthase genes). High activity of the promoter results in an increased transcription and translation of the promoter regulated fluorescent gene. This results in fluorescent protein accumulation and increases the overall fluorescence of the floating biofilm. Therefore, the measured floating biofilm fluorescence correlates with the cumulative promoter transcriptional activity of the entire bacterial population. As predicted, the cumulative P_*srfA*_ promoter activity in a Δ*comQ* floating biofilm, is lowered (Fig. 1 A). There is a short time window however (10 h – 30 h), where P_*srfA*_ is activated independently of the ComQXPA QS system. This indicates, that P_*srfA*_ activity is not completely abolished in the Δ*comQ* mutant. To support our claim further, we also semi-quantified surfactant production by the drop method (Fig. 1 B). Surfactants strongly decrease water surface tension, effectively increasing the water-solid interphase contact surface in a concentration dependent manner. The Δ*comQ* floating biofilm spent medium had a higher surface tension relative to the wild type (Fig. 1 B). This, alongside with the results shown on Fig. 1A confirm, that the production of surfactants is indeed decreased in the Δ*comQ* floating biofilm during growth and relative to the wild type.

**Figure 1:**
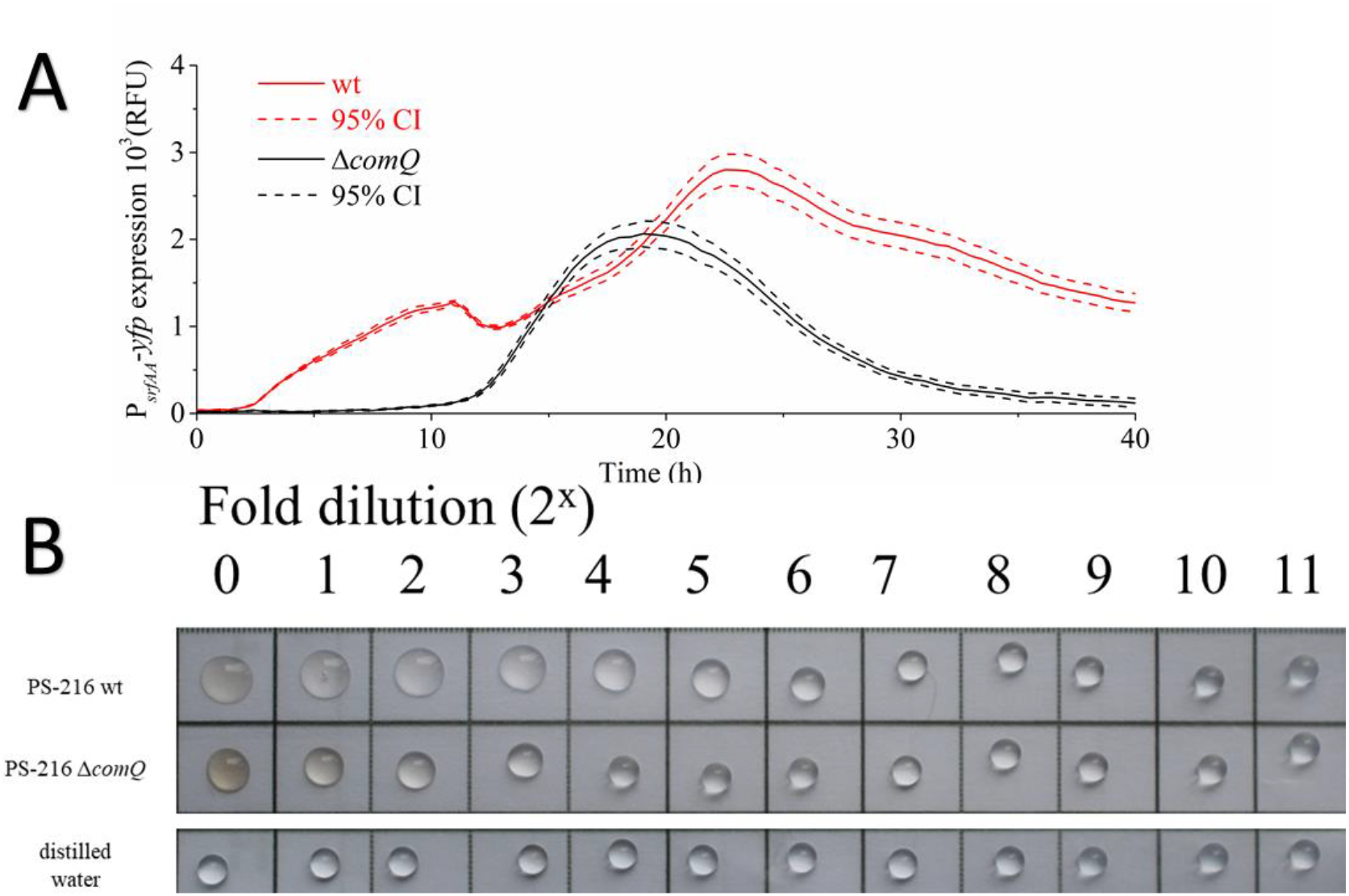
A) Floating biofilm fluorescence intensity of the P_*srfAA*_-*yfp* transcriptional reporter in the wild type (wt) and the QS mutant (Δ*comQ*) phenotype strain during static growth in MSgg medium at 37 °C. B) Semi quantification of surfactant concentrations in the wild type and QS mutant floating biofilm spent media with the droplet surface wetting assay after 40 hours of static growth in MSgg medium at 37 °C.

Additionally, the same difference is also evident in the NCIB 3610 strain, albeit it is much less dramatic (Fig. S1). NCIB 3610 appears to produce less surfactin compared to the PS-216 strain in general. To further validate our results we also show, that the spent media of a surfactin mutant does not significantly alter the surface tension of the spotted spent media droplet relative to the distilled water droplet (Fig. S1). Therefore we concluded, that the first part of our hypothesis is true, meaning that surfactin production is indeed diminished in the the PS-216 floating biofilm.

We then proceeded to check the phenotype of the floating biofilms in the wild type and the QS mutant by visually comparing both biofilms. In contrast with our prediction, we found that the Δ*comQ* strain forms a more robust floating biofilm (Fig. 2).

**Figure 2:**
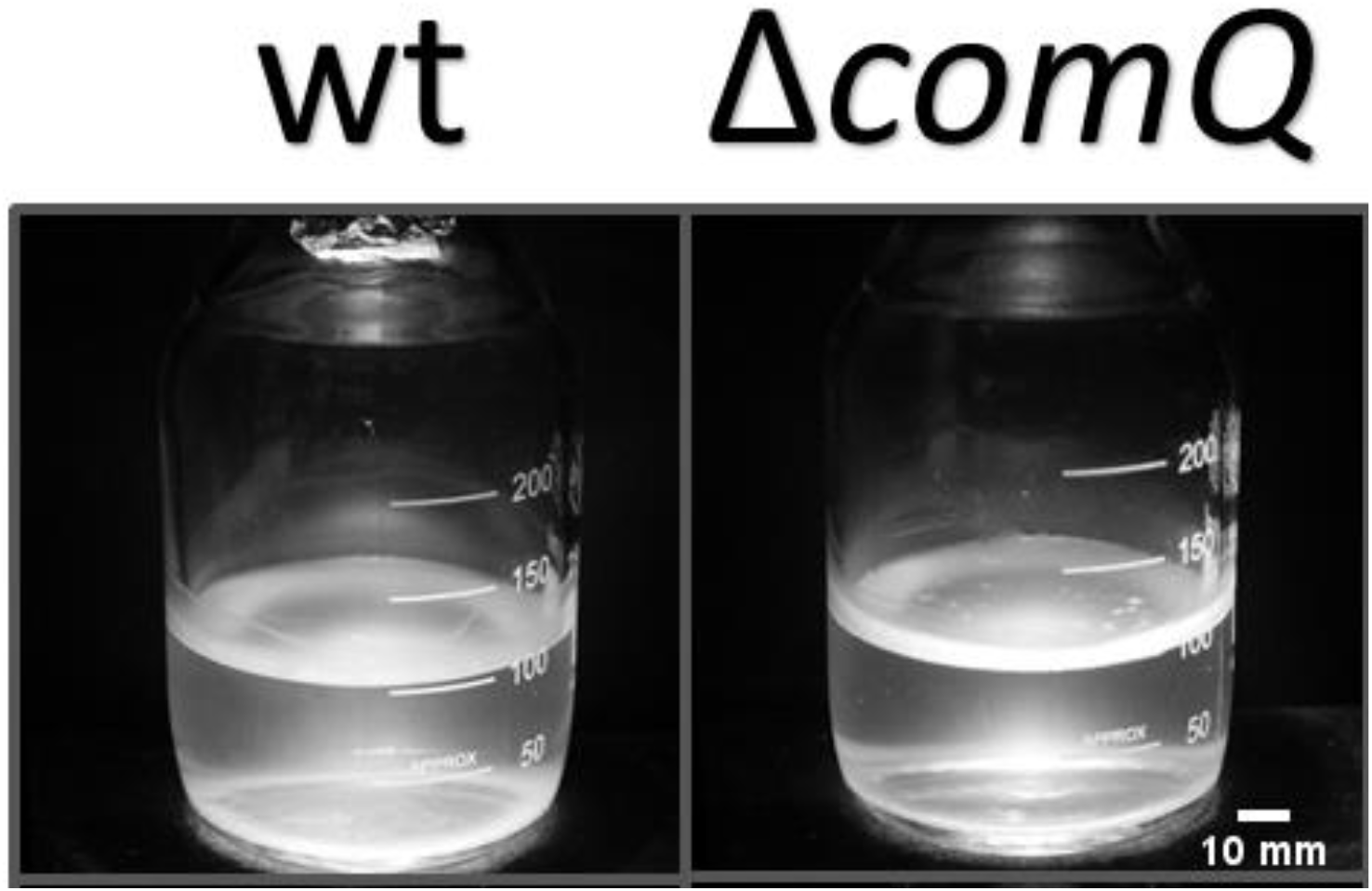
Floating biofilms of the *Bacillus subtilis* wild type PS-216 and the QS mutant (PS-216 Δ*comQ*) strains grown statically in MSgg medium at 37 °C and in 250 ml bottles for 16 hours.

Confocal microscopy of cells carrying a P_43_-*mKate2* transcriptional reporter construct, showed that in Δ*comQ* floating biofilms, cells were arranged in a thicker biofilm (up to 30 µm) than the wild type (up to 12 µm) strain (see the sides of the orthogonal display for biofilm thickness; Fig. 3). P_43_ is a constitutively active promoter (54), which enables the visualization of all metabolically active cells in the floating biofilm. The Δ*comQ* cells are less densely packed than those in the wild type floating biofilm and appear to be consisted mainly by chained cells (Figure 3), suggesting fast division. This difference was lost at 40 h, where only few fluorescing cells could be detected in both, wild type and Δ*comQ*, floating biofilms. Admittedly however, cells in a wild type biofilm are densely packed, whereas they appear to be more spread out in space in a Δ*comQ* floating biofilm.

**Figure 3:**
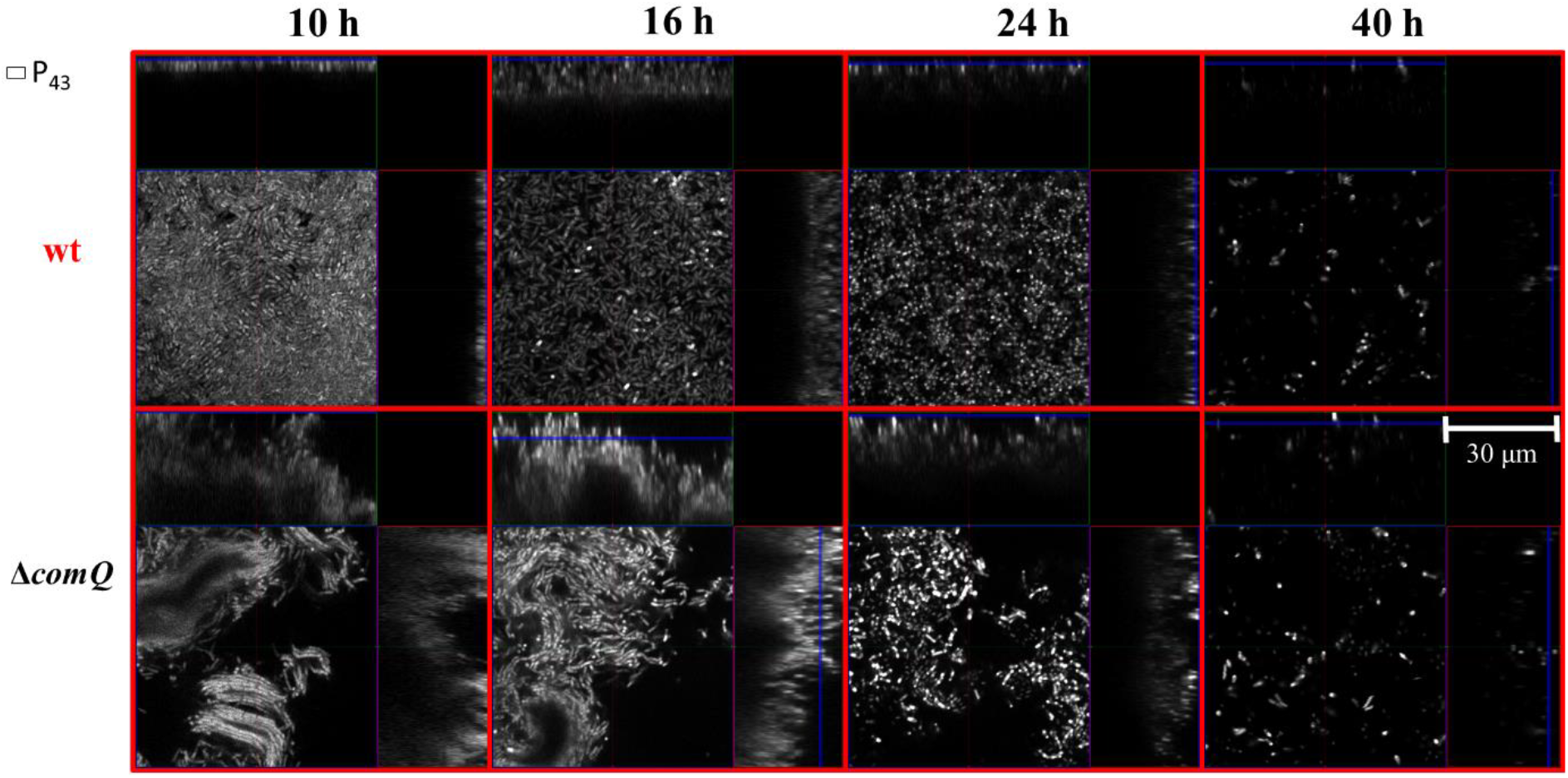
Floating biofilms of the PS-216 *Bacillus subtilis* wild type (wt) and the QS mutant (Δ*comQ*) phenotypes carrying a P_*43*_-*mKate2* transcriptional reporter during static growth in MSgg medium at 37 °C and visualized under a confocal microscope at different time points.

We also assayed hydrophobicity by following the diffusion of the methylene blue droplet placed on the top of the floating biofilm over time. The dye was slower to spread on the Δ*comQ* floating biofilm (Fig. S2). The Δ*comQ* floating biofilm was also rougher compared to the wild type (Fig. S3). As published work analyzing biofilm formation has been mostly performed using *B. subtilis* isolate NCIB 3610, we also compared floating biofilms formed by the NCIB 3610 wild type and its Δ*comQ* mutant. The difference in floating biofilm morphology between the two strains was less dramatic than for the PS-216 strains, but again the morphology of the Δ*comQ* mutant did not support the initial assumption that QS promotes floating biofilm formation (Fig. S3).

P_*epsA*_ and P_*tapA*_ control the transcription of the operons responsible for the synthesis of the major biofilm matrix polysaccharide and protein component, respectively. Both fluorescent transcriptional reporter constructs indicated a dramatically higher cumulative promoter activity in the Δ*comQ* compared to wild type floating biofilms (Fig. 4 A and C). This activity should therefore coincide with the floating biofilm sugar and protein content. The overall pellicle extract sugar content was higher in the extracellular polymer extracts of the Δ*comQ* floating biofilms compared to the wild type extracts (Student’s t-test and one way Mann-Whitney U test; P<0.05) at 16 h and 24 h of incubation (Fig. 4 B). The Δ*comQ* floating biofilm extracellular polymer extracts also contained a higher amount of protein (Student’s t-test and one way Mann Whitney U test; P < 0.05) relative to the wild type extracts at 16 h and 24 h of incubation (Fig. 4 D). This difference was lost at 40 h (Fig. 4 B and D), probably due to floating biofilm deterioration.

**Figure 4:**
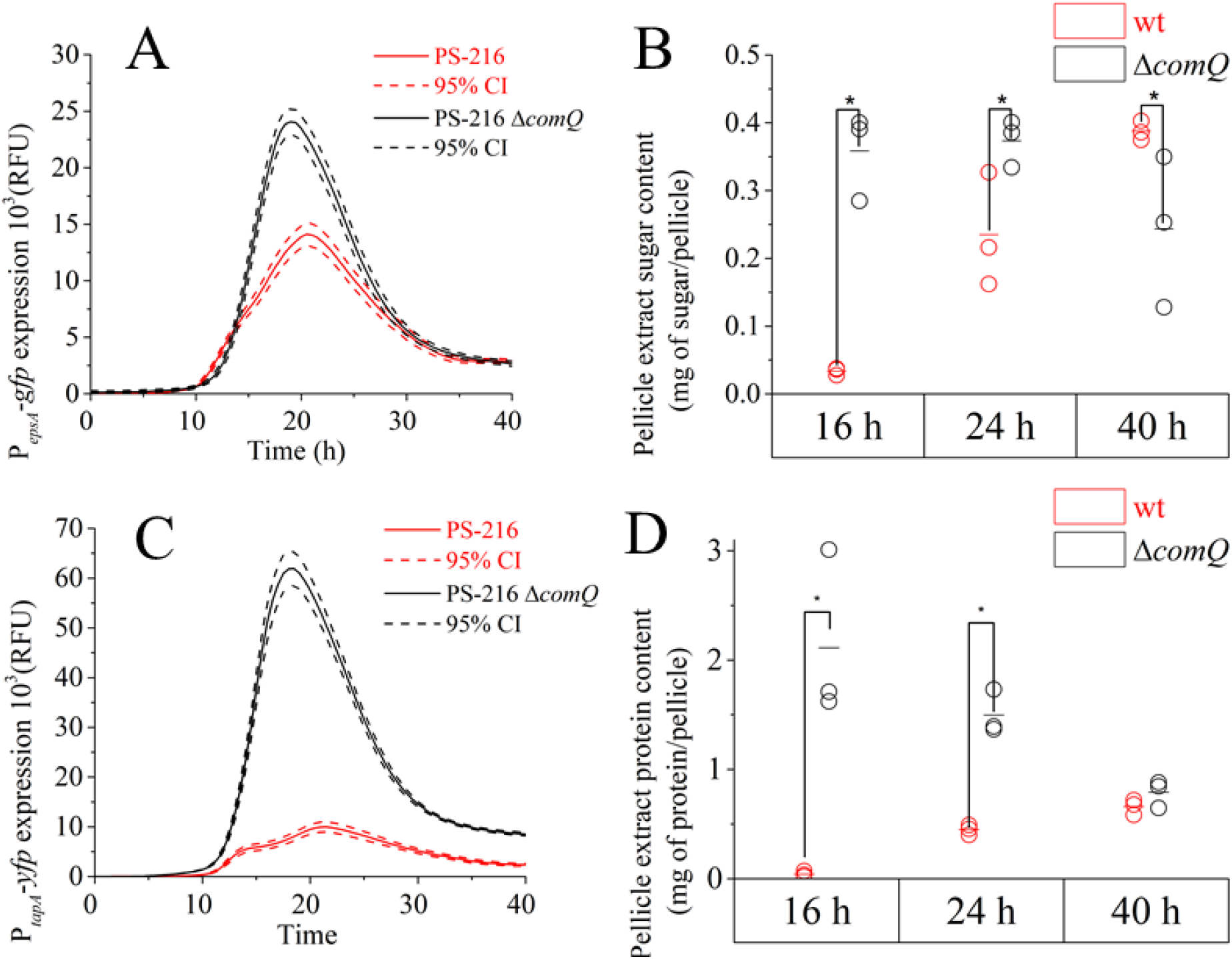
Floating biofilm fluorescence intensity, sugar, and protein quantity in *B. subtilis* PS-216 wild type (wt) strain and QS mutant (Δ*comQ)* during static growth in MSgg medium at 37 °C. A) Floating biofilm fluorescence intensity of the P_*epsA*_-*gfp* reporter constructs. B) Floating biofilm extract sugar content determined with the phenol–sulfuric acid method. C) Floating biofilm fluorescence intensity of the P_*tapA*_-*yfp* reporter constructs. D) Floating biofilm extract protein content determined with the Bradford assay. Each spot on the panels B and D represents the value of a biological repeat, with a horizontal dash showing the average value of all three measurements. Statistical significance was determined using a one-way Mann-Whitney test and a Student’s t-test (P<0.05).

Moreover, the cumulative P_*epsA*_ promoter activity in the NCIB 3610 wild type floating biofilm was also lower compared to the NCIB 3610 Δ*comQ* floating biofilm (Fig. S4). These results indicate that the findings above are not strain specific for the PS-216 strain.

Overall, these results refute our first hypothesis that the ComQXPA QS system promotes biofilm formation, despite increasing surfactin production.

### 1.4.2 Hypothesis 2

The ComQXPA QS system negatively regulates transcription of matrix producing genes (*epsA-O, tapA*) in the PS-216 *B. subtilis* strain at the single cell level.

QS Inactivation might promote floating biofilm formation by increasing the transcription and production of the biofilm matrix genes in each individual cell.

To test this, we normalized the cumulative P_*epsA*_ and P_*tapA*_ transcriptional activity to the P_43_ transcriptional activity. P_43_ is a strong, constitutively expressed promoter (54). By doing this, we obtained a simple quantitative measure of promoter transcriptional activity in single cells with an active P_43_ promoter. Results show that the normalized P_*epsA*_ promoter activity is actually higher in wild type cells and decreases in Δ*comQ* cells (Fig. 5A), despite the observation, that wild type forms an overall weaker floating biofilm (Fig. 1). Confocal micrographs of floating biofilms, formed by cells carrying the same transcriptional reporter constructs further confirm this result (Fig. S5). It appears that cells that do transcribe from the P_*epsA*_ promoter, do so less intensely in the Δ*comQ* strains (Fig. S5 B) despite the fact that cumulatively speaking (in terms of biofilm productivity) they produce more P_*epsA*_ regulated transcript (Fig. 4A). Normalized P_*tapA*_ promoter activity is similar in both cases (Fig. 5B). Therefore, on (Fig. 5), we can see that our estimates for single cell promoter activity for P_*epsA*_ and P_*tapA*_ do not support the prediction made in Hypothesis 2. This indicates that QS regulation of biofilm matrix component gene transcription is not inhibitory, and could not explain the difference in floating biofilm formation rates, because P_43_ normalized promoter activity of P_*epsA*_ and P_*tapA*_ is not significantly lower in wild type floating biofilms. We further reasoned, that a bacterial population with an inactive QS forms floating biofilms at a faster rate mostly due to an increased number of metabolically active cells in the population, providing an explanation for the second hypothesis.

**Figure 5:**
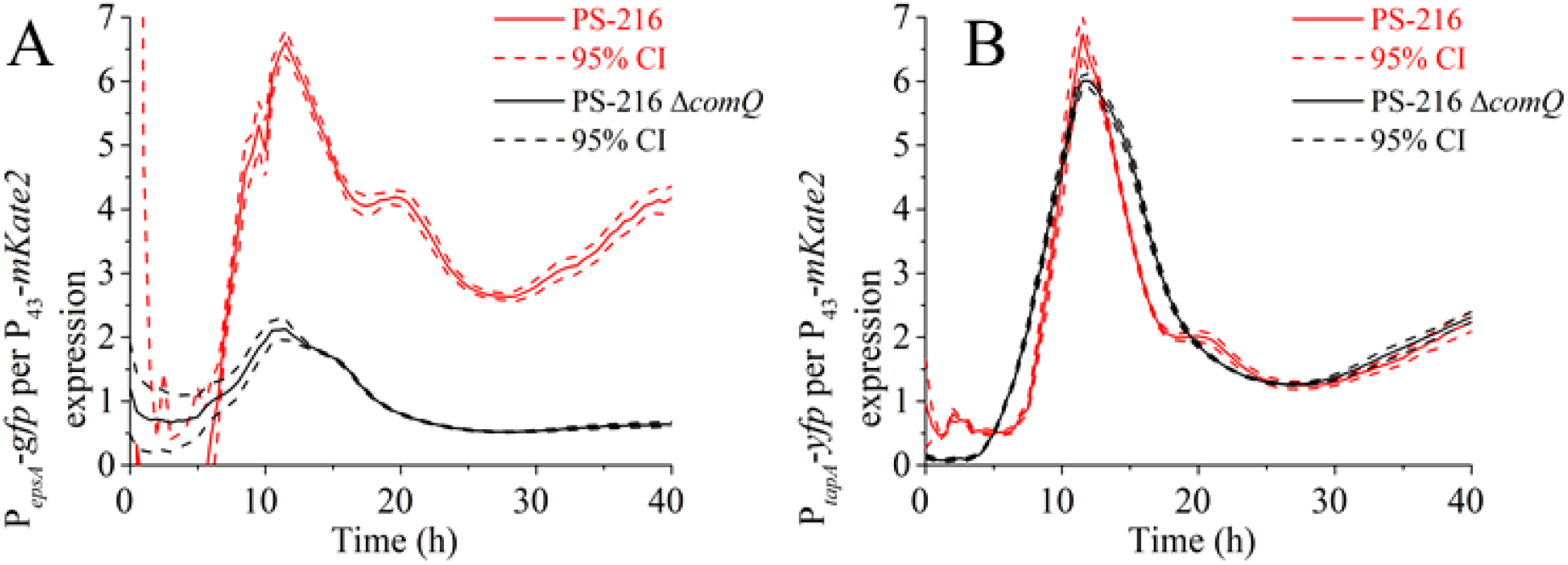
A) Floating biofilm fluorescence intensity of the P_*epsA*_-*gfp* normalized per P_*43*_-*mKate2* fluorescence and B) floating biofilm fluorescence intensity of the P_*tapA*_-*yfp* fluorescence normalized per P_*43*_-*mKate2* fluorescence in *Bacillus subtilis* PS-216 wild type strain and QS mutant during static growth in MSgg medium at 37 °C.

### 1.4.3 Hypothesis 3

A developing wild type floating biofilm contains more cells committed to sporulation during biofilm development compared to the Δ*comQ*

Our final hypothesis assumed that the QS system in floating biofilms slows down growth rates and increases stochastic sporulation initiation. Therefore, even if single cell biofilm matrix production is greater in wild type cells (e.g. P_43_ normalized P_*epsA*_ promoter activity on Fig. 5A), or equal (e.g. P_43_ normalized P_*tapA*_ promoter activity on Fig. 5B) a QS inactivated mutant cell population will produce more biofilm matrix nevertheless, because it has a greater number of metabolically active cells at any given time point.

To test this hypothesis, we measured the colony forming units (CFU) in the wild type and Δ*comQ* floating biofilms (Fig. 6A). Additionally we also measured the quantity of heat resistant CFU in the same floating biofilms, to determine the fraction of endospore forming cells in a floating biofilm (Fig. 6 B). Both results were in accord with our hypothesis, showing not only that a wild type floating biofilms has fewer CFU at 16 h and 24 h (Fig. 6A), but also that a significantly higher fraction of those wild type cells are actually spores (Fig. 6B). Both phenotypes however end up with a similar CFU count after 40 h, and all of the CFU are heat resistant. We also measured expression of the P_*spoIIQ*_ (the promoter expressed during early stages of endospore development) and normalized its activity per P_*43*_ activity. We found the P_*spoIIQ*_ promoter activity being higher in the wild type than in the Δ*comQ* floating biofilm. Also, addition of the exogenous ComX restored the P_*spoIIQ*_ activity in the QS deficient mutant to wild type levels (Fig. 6 D).

**Figure 6:**
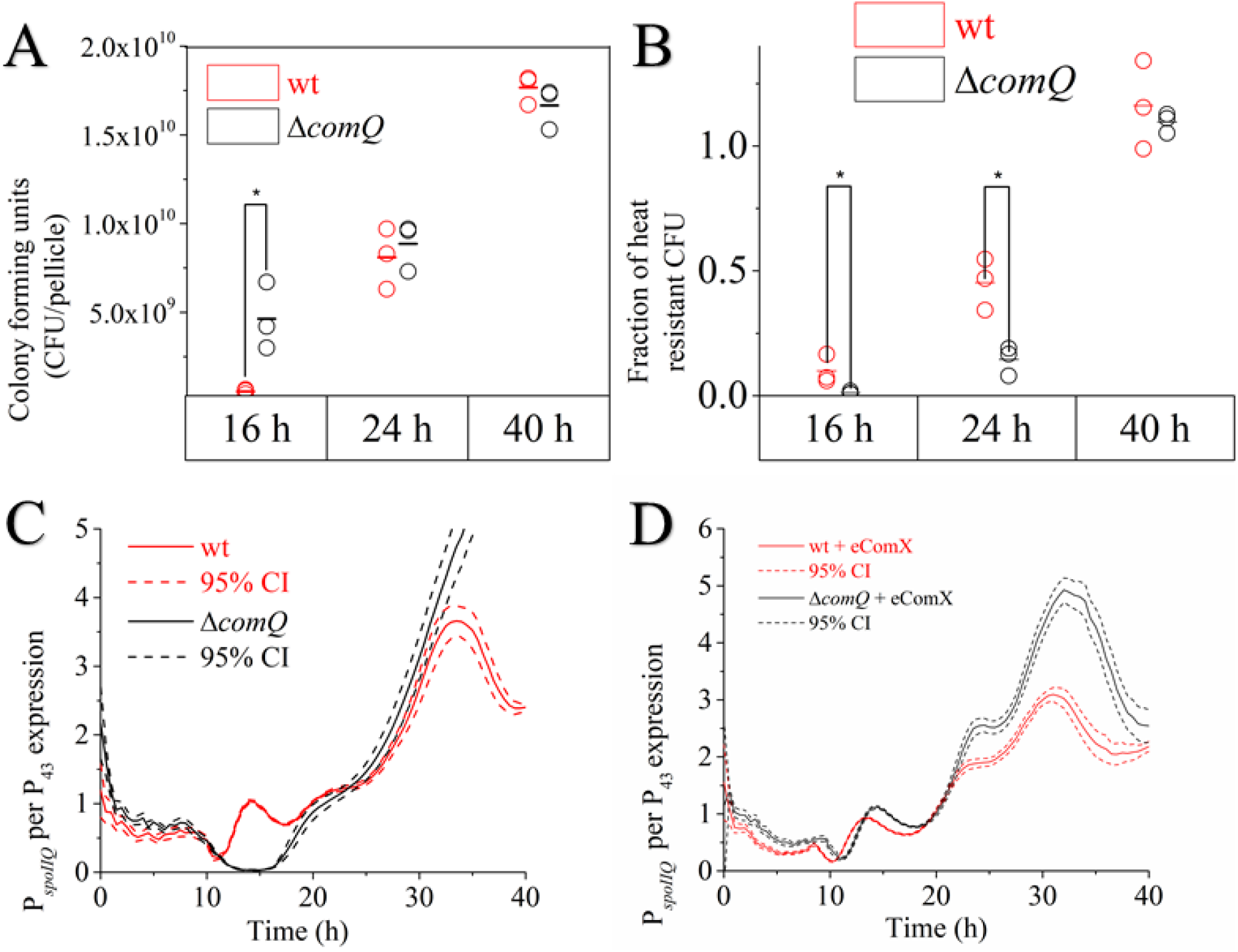
Comparisons of *B. subtilis* PS-216 wild type (wt) and QS mutant (Δ*comQ*) during static growth in MSgg medium at 37 °C. A) Total floating biofilm CFU counts, B) Floating biofilm heat resistant CFU fraction, C) Floating biofilm P_*spoIIQ*_-*yfp* fluorescence normalized per P_*43*_-mKate2 fluorescence, D) Floating biofilm P_*spoIIQ*_-*yfp* fluorescence normalized per P_*43*_-mKate2 fluorescence with exogenous heterologously expressed ComX. Each spot on the panels A and B represents the value of a biological repeat, with a horizontal dash showing the average value of all three measurements. Statistical significance was determined using a one-way Mann-Whitney test and a Student’s t-test (P<0.05).

The difference in timing of the P_*spoIIQ*_ promoter activity between the wild type and Δ*comQ* floating biofilm was also observed by confocal microscopy (Fig. 7), and suggests that the wild type forms pre-spores already after 10 h of incubation. In contrast no pre-spores are evident in the Δ*comQ* floating biofilm at this time. This difference is apparent in spite of the faster growth rates of the Δ*comQ* mutant (Fig. 6A). On the same confocal micrographs we can also observe individual pre-spores within *B. subtilis* cells (Fig. 7), because the P_*spoIIQ*_ regulated fluorescent product is highly localized in the pre-spore (55).

**Figure 7:**
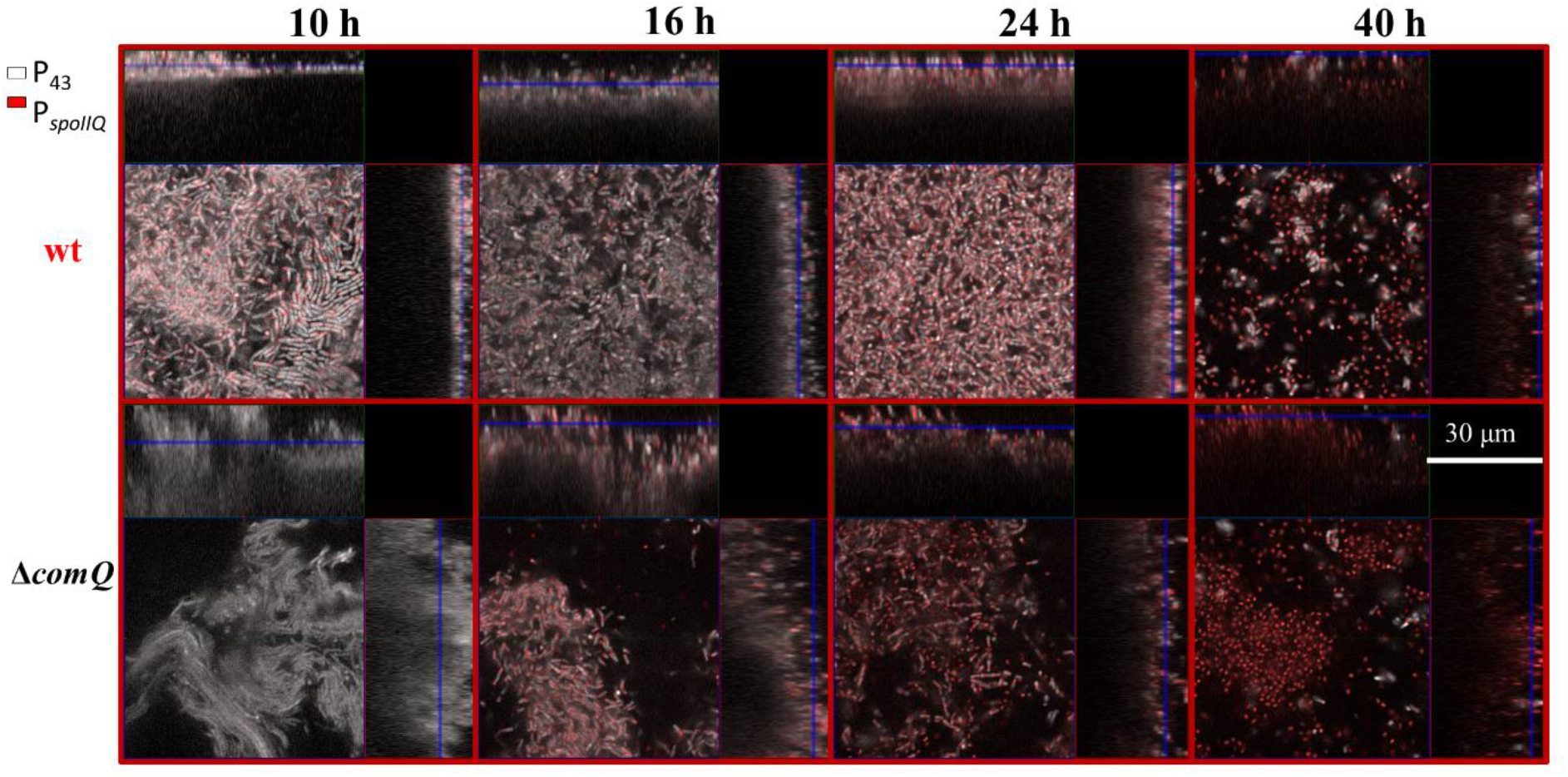
Confocal visualizations of P_*spoIIQ*_-yfp (false colored red) and P_*43*_-*mKate2* (false colored white) expression in wild type (wt) and QS mutant (Δ*comQ*) floating biofilms during static growth in MSgg medium at 37 °C.

Collectively, these results indicate that the ComQXPA QS system operates as a control switch device that limits the investment made into population growth by promoting commitment to late growth adaptive processes, like sporulation. In the mutant that lacks *comQ*, this QS dependent brake, which enables bet hedging behavior, is released and the fitness of the population increases transiently leading to promoted floating biofilm formation.

## 1.5 DISCUSSION

The ComQXPA QS system upregulates surfactin production (21, 56, 57) and because surfactin promotes transcription of the *epsA-O* operon (49) it has been often assumed that ComX dependent QS and/or surfactin promote biofilm formation (4, 12, 13, 16, 39, 51–53). In this work, we further test this assumption and confirm that ComX does indeed appear to positively regulate transcription of the *epsA-O* operon in single cells (Fig. 5) as previously reported (49). However, we further show, that the ComX deficient mutant (Δ*comQ*) forms thicker floating biofilms nonetheless, with more cells and matrix, but less spores during the early stages of biofilm formation. Our results suggest that this is because ComX acts as a switch that promotes early commitment to sporulation in a sub-population of cells and thus contributes to bet hedging in *B. subtilis*.

Our results therefore re-confirm (Fig. 1 and Fig. 5A) that the ComQXPA QS system increases surfactin production (21) and the transcription of the *epsA-O* operon per cell (49). However we could not conclude that QS promotes overall formation of floating biofilms (Fig. 2), as predicted in hypothesis 1. While P_43_ normalized P_epsA_ promoter activity appeared to be positively regulated by ComX the P_43_ normalized P_*tapA*_ promoter activity was comparable in both the wild type and Δ*comQ* biofilms (Fig. 5). This expression pattern is consistent with recent reports that P*epsA* and P*tap*A are not co-regulated (59). However, expression pattern of both genes in the △*comQ* biofilm does not explain the thicker biofilms of the mutant where much higher P_43_ normalized activity of both the P_*epsA*_ and P_*tapA*_ promoters in the Δ*comQ* mutant would be expected if this was the case (Fig. 5). We therefore rejected the second hypothesis.

The confocal micrographs indicate that the wild type cells, which are more segmented at 10h, show high *epsA-O* transcriptional activity at this time (Fig. S5), as opposed to the QS mutant cells, which are more chained and have increased *tapA* transcription (Fig. S6). This is consistent with recent report by van Gestel and co-workers (59) who associate high *epsA-O* transcription with segmented cells and high *tapA* transcription with chained cells (58).

Biofilms have been mostly studied using either the 168 domesticated laboratory strain or the NCIB 3610 undomesticated strain (17). We opted to conduct our research on the PS-216 strain, which is very similar to both before mentioned strains in terms of genomic sequence (17, 59). Both strains, the NCIB 3610 and the PS-216 are undomesticated, however, the NCIB 3610 strain has a drastically lower surfactin production (Fig. S1) and forms weaker floating biofilms (Fig. S3) than PS-216. Although not as dramatic as in PS-216, the NCIB 3610 QS mutant still shows a higher cumulative transcription level of the *epsA-O* operon compared to the wild type NCIB 3610 (Fig. S4). We therefore argue, that our findings are generally applicable to *B. subtilis* species as opposed to being PS-216 specific.

### 1.5.1 ComX represents an important switch towards bet hedging behavior

Phenotypic plasticity is defined as the ability of an individual (in our case a bacterial population) to adapt to a changing environment (60). The phosphorelay is interpreted as being a noise generation interface, which enables bi-stable sporulation of a sub-population of cells attaining high enough levels of Spo0A-P (61). This is also believed to enable bet hedging behavior (14, 62, 63). Therefore, the phosphorelay affecting bi-stable expression of operons in individual cells, confers greater phenotypic plasticity to the bacterial population. *Bacillus subtilis* is known to initiate stochastic sporulation independently from responsive triggers (e.g. nutrient starvation) in order to “bet hedge” (14). Our results also corroborate this observation in floating biofilms, with bet hedging being most apparent in terms of heat resistant CFU (Fig. 6). While at the 10 h time point the wild type strain has already committed a part of the population to sporulation. The QS mutant, despite growing faster than the wild type, enters this developmental process more uniformly at the later time, and thus avoiding bet hedging (Figure 7).

### 1.5.2 The interplay of external stimuli and auto-inducing signaling in the molecular regulation of biofilm formation

The phosphorylation state of ComA is also affected independently from ComX via the PhrC (CSF) signalling molecule and the RapC phosphatase (64). CSF is also considered as being a quorum sensing molecule, however, ComX has a greater impact on the ComA phosphorylation state and surfactin transcription (65). In this work, therefore, we limited ourselves to study mostly the effect of the ComQXPA QS system on floating biofilm formation. Additionally, there are several other Rap-Phr signal/phosphatase pairs, which can affect the phosphorelay system (66). Interestingly, when the ComQXPA QS system was inactivated, sporulation appeared to be less stochastic, thus suggesting also that the ComQXPA QS system is the main driver of stohasticity in *B. subtilis*. This could explain the apparent redundancy of the two systems, both regulating ComA phosphorylation. The Rap/Phr might contribute to a starvation driven intiation of sporulation, while the ComQXPA QS system might contribute to stochastic sporulation initation, but this remains to be investigated.

### 1.5.3 Fitness benefits of public goods and the role of cannibalism

Bacterial public goods are defined as being accessible to all cells in a population and that they provide a fitness benefit (67). Firstly, in regards of accessibility, it has been already shown that due to spatial segregation, the production of extracellular polysaccharides cannot be exploited against in a biofilm (68, 69). As extracellular polysaccharides (Eps) in the biofilm matrix are are not accessible to every cell in the population they may be considered a private rather than public good (67). Secondly, the fitness benefit is often very context dependent, and also differs in regards to the selective pressure that is being exerted (70). Our results show, that in the end in terms of fitness, defined as an increase in the copies of an allele (71), neither the wild type nor the QS mutant have a significant advantage under the tested set conditions at the 40 h time point (Fig. 6 A and B), since both the wild type and QS mutant end up with roughly comparable allele copy number or spores. Therefore, the environment is either non-selective or the QS system (and the phenotype changes it affects) cannot be considered beneficial. However, we also observed a stark difference in growth rates, sporulation efficiency and biofilm matrix component production at earlier time points, meaning that several bottlenecks could be construed with these differences in mind. Those could confer a fitness advantage to either the wild type or the QS mutant strain, but for this research to be sensible, we would have to at least argue as to why we think that these bottlenecks are present in *B. subtilis*’ natural environment. The soil, which is arguably the natural habitat of *B. subtilis*, is often exposed to droughts at noon, when the sun is at its highest. Additionally, nutrients in the environment might also periodically get depleted and make conditions where sporulation confers a fitness advantage. Therefore, a bet hedging strategy would most likely be beneficial. We here show that QS mutant does not assume this bet hedging strategy in regard to spore formation during early stages of biofilm development (Fig. 6 B and 7). This suggests that the wild type strain may have a fitness advantage over the QS mutant if selective pressure (in form of a sudden temperature increase, drought or nutrient depletion) appeared during growth (14, 15, 63).

Our results also show that the wild type strain grows more slowly than the QS mutant strain. This might, on the other hand, represent a disadvantage in an environment, where fast nutrient utilization and overgrowth is required in order to outcompete other bacteria which occupy a similar ecological niche. Indeed, our results indicate that the QS mutant grows at faster rates in static cultures (Fig. 6 A) and in shaken cultures (23). Therefore, it would be expected that QS mutant would most likely have an advantage over the competitor strains if selective pressure would operate at the level of growth rates (72). On the other hand, a more slowly growing wild type phenotype strain might utilize scarce nutrients more efficiently and also through bet hedging mitigate the consequences of incorrectly predicting a non-favorable environmental fluctuation. One of the reasons why a QS mutant won’t overgrow the wild type strain in nature is possibly due to the private link between signal and response in the *B. subtilis* QS system, described in (23).

### 1.5.4 Floating biofilm formation is stunted due to bet hedging

*Bacillus subtilis* is widely known for its bet hedging behavior in terms of spore formation, where a sub-population of cells initiates sporulation stochastically, regardless of the sensed external environmental stimuli (e.g. nutrient starvation) (14). In a wild type population, the endospore forming process started already at the onset of floating biofilm formation. The bacterial population with an inactivated QS however, did not exhibit such behavior. So to sum our findings up, we showed that the ComQXPA QS system serves as a switch that down modulates investment into growth and assures early investment into sporulation. This, on one hand, enables bet hedging behavior, but on the other hand, burdens cells with the metabolic costs of early sporulation initiation and ultimately slows down the floating biofilm formation process.

## 1.6 MATERIALS AND METHODS

### 1.6.1 Bacterial strains and strain construction

All *Bacillus subtilis* strains used in this study are listed in Table 1. The recombinant strains were constructed by transforming specific markers into *B. subtilis* recipients. The recipient strains were grown in competence medium (CM) at 37°C (73) and after 6 h of incubation chromosomal or plasmid DNA was added to competent cells. Transformants were selected on LB agar with specific antibiotic (chloramphenicol (Cm) 10 μg/ml, kanamycin (Kan) 50 μg/ml or spectinomycin (Spec) 100 μg/ml) at 37°C. Competence in strains with *comQ* deletion was induced by the addition of exogenous ComX as previously described (24). Integration of P_*epsA*_-*gfp* reporter fusion into different strains was performed by transformation with YC164 genomic DNA (41) (Table 1). Integration of *srfA::Tn917* (mls) mutation into different strains was performed by transformation with BM1044 genomic DNA (74) (Table 1). Strains with P_*tapA*_-*yfp* reporter fusion were constructed by transformation with BM1115 genomic DNA (75). To prepare strains carrying the P_*spoIIQ*_-*yfp* reporter fusion the plasmid pKM3 (55) was transformed into indicated recipients (Table 1). To transform strains with the P_*43*_-*mKate2* reporter fusion (Table 1), the plasmid pMS17 with the kanamycin resistance marker and the plasmid pMS7 with the chloramphenicol resistance marker were used (Table 3). To prepare strains with the P_*srfAA*_-*yfp* reporter fusion DL722 chromosomal DNA was transformed into different strains (50). To construct the NCIB 3610 QS mutant the plasmid pMiniMAD2-updowncomQ (Table 3) was used (24). The plasmid was transformed into *B. subtilis* NCIB 3610 *comI*^*Q12L*^ as previously described (24, 76).

**Table 1:** Bacterial strains used in this study.

| Strain name | Background | Genome description | Reference |
| --- | --- | --- | --- |
| ***Bacillus subtilis* strains** |  |  |  |
|  | PS-216 | wt | (79) |
| BM1127 | PS-216 | $\Delta comQ$ | (24) |
| BM1629 | PS-216 | *sacA::P<sub>43</sub>-mKate2* (Cm) | this work |
| BM1630 | PS-216 | $\Delta comQ$<br>*sacA::P<sub>43</sub>-mKate2* (Cm) | this work |
| BM1625 | PS-216 | *amyE::P<sub>spoIIQ</sub>-yfp* (Sp)<br>*sacA::P<sub>43</sub>-mKate2* (Cm) | this work |
| BM1626 | PS-216 | $\Delta comQ$<br>*amyE::P<sub>spoIIQ</sub>-yfp* (Sp)<br>*sacA::P<sub>43</sub>-mKate2* (Cm) | this work |
| BM1613 | PS-216 | *amyE::P<sub>tapA</sub>-yfp* (Sp)<br>*sacA::P<sub>43</sub>-mKate2* (Cm) | this work |
| BM1614 | PS-216 | $\Delta comQ$<br>*amyE::P<sub>tapA</sub>-yfp* (Sp)<br>*sacA::P<sub>43</sub>-mKate2* (Cm) | this work |
| BM1629 | PS-216 | *sacA::P<sub>43</sub>-mKate2* (Kan) | this work |
| BM1630 | PS-216 | $\Delta comQ$<br>*sacA::P<sub>43</sub>-mKate2* (Kan) | this work |
| BM1631 | PS-216 | *amyE::P<sub>epsA</sub>-gfp* (Cm)<br>*sacA::P<sub>43</sub>-mKate2* (Kan) | this work |
| BM1622 | PS-216 | $\Delta comQ$<br>*amyE::P<sub>epsA</sub>-gfp* (Cm)<br>*sacA::P<sub>43</sub>-mKate2* (Kan) | this work |
| DK1042 | NCIB 3610 | *comI<sup>Q12L</sup>* | (80) |
| BM1667 | NCIB 3610 | $\Delta comQ$<br>*comI<sup>Q12L</sup>* | this work |
| BM1623 | NCIB 3610 | *comI<sup>Q12L</sup>* *amyE::P<sub>epsA</sub>-gfp* (Cm) | this work |
| BM1624 | NCIB 3610 | *comI*<sup>Q12L</sup><br>*ΔcomQ*<br>*amyE::P<sub>epsA</sub>-gfp* (Cm) | this work |
| YC164 | NCIB 3610 | *amyE::P<sub>epsA</sub>-gfp* (Cm) | (41) |
| BM1115 | PS-216 | *amyE::P<sub>tapA</sub>-yfp* (Sp) | (75) |
| BM1097 | PS-216 | *amyE::P<sub>hyperspank-mKate2</sub>* (Cm) | (75) |
| BM1044 | PS-216 | *srfA::Tn917* (mls) | (74) |
| BM1673 | PS-216 | *ΔcomQ*<br>*srfA::Tn917* (mls) | this work |
| DL722 | 3610 | *amyE::P<sub>srfAA</sub>-yfp* (Sp) | (50) |
| BM1454 | PS-216 | *amyE::P<sub>srfAA</sub>-yfp* (Sp) | this work |
| BM1455 | PS-216 | *ΔcomQ*<br>*amyE::P<sub>srfAA</sub>-yfp* (Sp) | this work |
| ***Escherichia coli*** |  |  |  |
| ED367 | BL-21 (DE3) | pET22(b) – *comQ comX* from *B. subtilis* 168 (Amp) | (21) |

The P_*43*_-*yfp* construct from Pkm3-p43-*yfp* plasmid (75) was digested with EcoRI and BamHI and ligated into the pSac-Cm (77) plasmid to form the pEM1071 plasmid carrying a P_*43*_-*yfp* fusion inside the *sacA* integration site. P_*43*_ is a constitutively expressed promoter in *Bacillus subtilis* (54), located upstream from the *yfp* gene in the pEM1071 plasmid. This vector was then digested with HindIII and BamHI restriction enzymes to remove *yfp* gene from the original vector and then simultaneously ligated with the digested *mKate2* fragment to construct the plasmid pMS7. The *mKate2* sequence was PCR amplified using the P3F and P3R primer pair (Table 2) and the genomic DNA of BM1097 as the template (78). To construct the plasmid pMS17, the plasmid pMS7 was digested with EcoRI and BamHI restriction enzymes and the digested P_*43*_-*mKate2* fragment was ligated into pSac-Kan EcoRI/BamHI restriction sites (77).

**Table 2:**
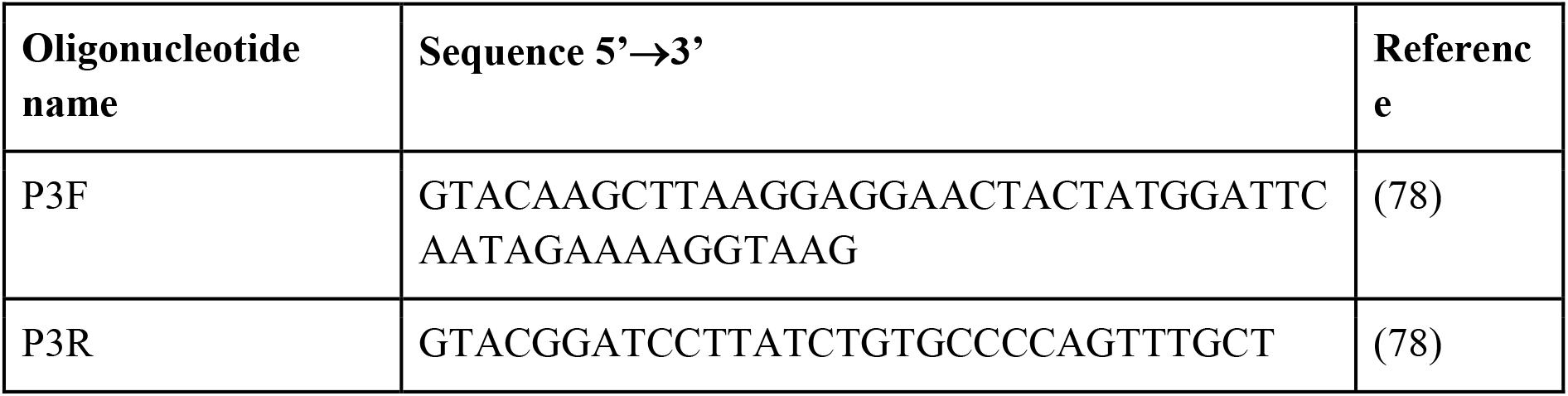
Oligonucleotides used in this study.

**Table 3:** Plasmids used in this study.

| Plasmid name | Description | Reference |
| --- | --- | --- |
| pKM3 | *amyE::P<sub>spoIIQ</sub>-yfp* (Sp, Amp) | (55) |
| Pkm3-p43-yfp | *amyE::P<sub>43</sub>-yfp* (Sp, Amp) | (55) |
| pSac-Kan | *sacA::kan* (Amp) | (77) |
| pSac-Cm | *sacA::cat* (Amp) | (77) |
| pEM1071 | *sacA::P<sub>43</sub>-yfp* (Cm, Amp) | this work |
| pMS7 | *sacA::P<sub>43</sub>-mKate2* (Cm, Amp) | this work |
| pMS17 | *sacA::P<sub>43</sub>-mKate2* (Kan, Amp) | this work |
| pMiniMAD2-updowncomQ | pMiniMAD2 with updown *comQ* between EcoRI and SalI sites (Mls, Amp) | (24) |

### 1.6.2 Growth conditions

Bacterial overnight cultures were grown in LB medium with the appropriate antibiotics at 37 °C with shaking at 200 rpm. Floating biofilms were grown by re-suspending an overnight culture (1 % v/v) in liquid MSgg medium (35) and incubating the culture in static conditions at 37 °C for up to 40 h. The heterologously expressed ComX in M9 spent medium was prepared as described previously (24).

### 1.6.3 Biochemical composition of extracellular polymers and CFU counts determination in floating biofilms

Floating biofilms were grown in 20 ml of liquid MSgg medium at 37 °C for up to 40 h in Petri dishes (90 mm diameter). At indicated times, floating biofilms were collected and transferred into two centrifuge tubes containing 1 ml of physiological saline solution. Floating biofilms were kept on ice during sonication with the MSE 150 Watt Ultrasonic Disintegrator Mk2 at three 5-seconds bursts and amplitude 15 μm. After floating biofilm disintegration cell counts were determined by colony forming units (CFU) on LB agar after 24 h of incubation at 37 °C. Spores were enumerated as CFUs/ml after heating cell suspensions to 80 °C for 30 minutes. The spore fraction was determined by dividing heat resistant CFUs with total CFUs. Extracellular polymers were extracted from floating biofilms as described previously (34). The total sugar content in the extracted polymer fraction was determined by the phenol–sulfuric acid method as described before (81) and the total protein content by the Bradford protein assay (82).

### 1.6.4 Spent media droplet surface wetting assay

Floating biofilm spent media were collected after 40 hours of static growth in MSgg medium at 37 °C. A two-fold dilution series in distilled water was made and 20 μL droplets of the spent media and all of the serial dilutions were gently placed on the top of a parafilm strip. The wetting surface area of the droplet corresponds with the droplet surfactant concentration. Greater surfactant concentrations will cause the droplet to cover a bigger parafilm surface area, thus appearing bigger from a top-down perspective. Distilled water droplets serve as a control. The *srfA* mutant spent medium dilutions indicate that the decrease in surface tension is mostly due to the synthesized surfactin and no other surfactant the wild type or QS mutant strain might produce, since they cover a similar surface than a distilled water droplet.

### 1.6.5 Expression of different transcriptional reporters during floating biofilm formation

Floating biofilms were grown in 200 µl MSgg medium in sterile 96-well black transparent bottom microtiter plate. If needed, and as indicated, strains were supplemented with ComX. Each strain was tested in three time independent replicates. Results are presented as averages of all the wells with 95 % confidence intervals. The lid was sealed with micropore tape and the microtiter plate was incubated in the Cytation 3 imaging reader (BioTek, United States) at 37 °C without shaking. Optical density at 650 nm and fluorescence intensity were measured in half hour intervals for up to 60 h. The gain settings, excitation and emission wavelengths for every transcriptional reporter are shown in Table 4.

**Table 4:**
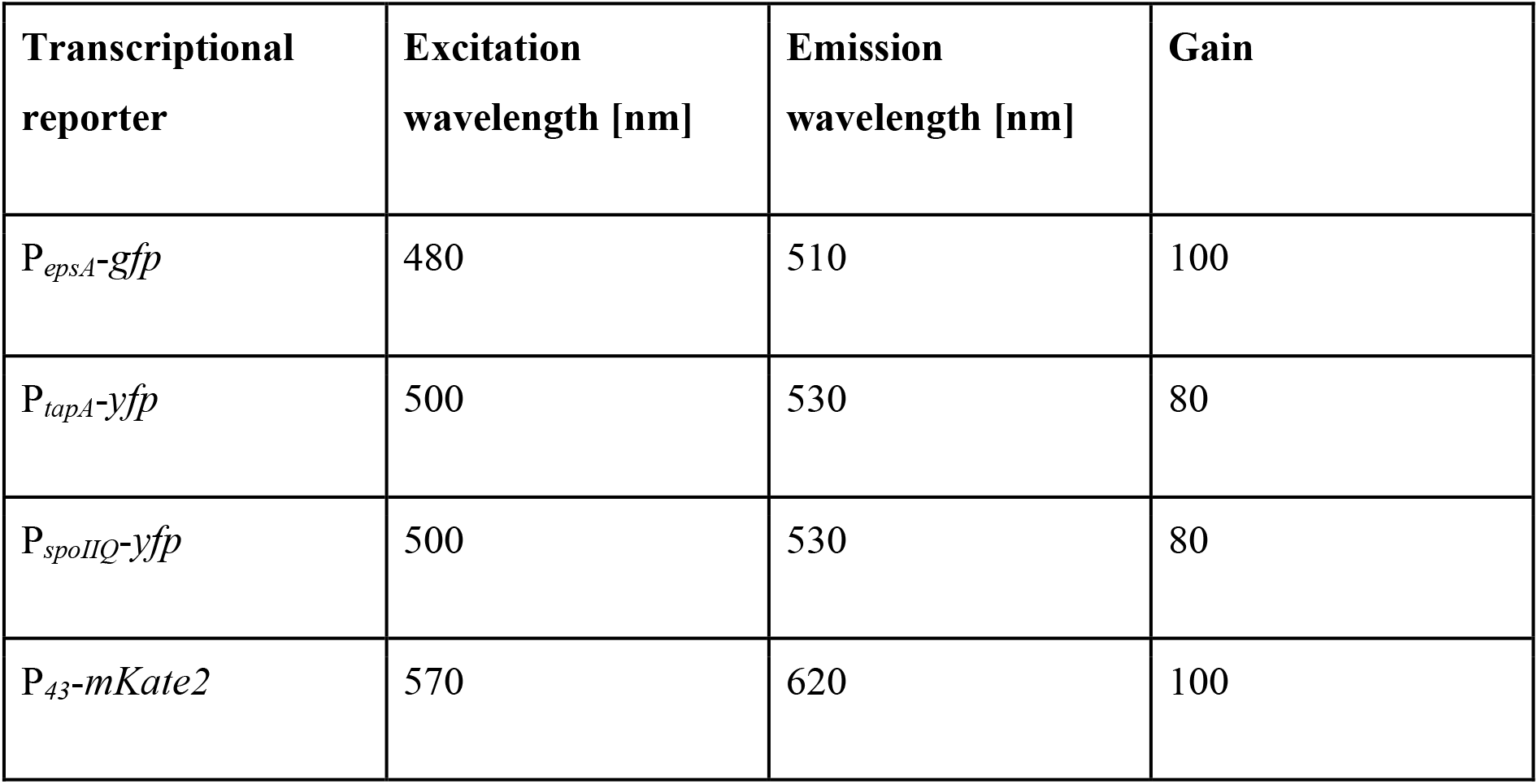
Emission and excitation wavelengths and gain settings for transcriptional reporters used in this study.

| Transcriptional reporter | Excitation wavelength [nm] | Emission wavelength [nm] | Gain |
| --- | --- | --- | --- |
| $P_{epsA}$-*gfp* | 480 | 510 | 100 |
| $P_{tapA}$-*yfp* | 500 | 530 | 80 |
| $P_{spoIIQ}$-*yfp* | 500 | 530 | 80 |
| $P_{43}$-*mKate2* | 570 | 620 | 100 |

Isogenic strains without the fluorescent reporter (background strains) were always cultured in the same microtiter plate as experimental strains. To calculate the final expression of the experimental strain the average fluorescence intensity of its background strain (grown in 8 wells) was deducted from the fluorescence intensity of the experimental strain at each time point to subtract an estimation of background fluorescence.

### 1.6.6 Floating biofilm morphology, hydrophobicity estimation and confocal laser scanning microscopy

Floating biofilms were grown in 2 ml of MSgg medium in 35-mm diameter glass bottom Petri dishes and their visual appearance documented by taking photos of images obtained with the Leica WILD M10 stereomicroscope. Floating biofilm hydrophobicity was evaluated by spotting a 50 μl droplet of 0.5 % (w/v) methylene blue solution in the center of the floating biofilm and the diffusion of the dye recorded over time by a video camera and then edited by Lightworks 14.5 trial video editing software. Cells in floating biofilms were visualized by confocal laser scanning microscope after removing the spent medium from underneath the floating biofilm by careful pipetting. We used the 100x immersion objective (NA 0.4) and roughly determined the bottom of the floating biofilm to set up a Z-stack. Floating biofilms carrying transcriptional reporters fused to specific promoters or/and constitutive promoters were then visualized using an LSM 800 microscope (Zeiss, Germany). Excitation laser wavelength, emission filters setup and other laser settings are shown in Table 5. Averaging was set on 8, scanning was unidirectional, pixel time equaled 0.58 μs and the frame time was 9.11 s.

**Table 5:** Confocal laser scanning microscope settings for different transcriptional reporters used in this study.

| Transcriptional reporter | Excitation Laser wavelength [nm] | Emission filter [nm] | Laser power [%] | Master gain [v] | Pinhole [$\mu\text{m}$] |
| --- | --- | --- | --- | --- | --- |
| **BM1631 and BM1622** |  |  |  |  |  |
| $P_{epsA}\text{-}gfp$ | **488** | **400-580** | **15** | **650** | **71** |
| $P_{43}\text{-}mKate2$ | **561** | **585-700** | **2.5** | **600** | **84** |
| **BM1613 and BM1614** |  |  |  |  |  |
| $P_{tapA}\text{-}yfp$ | **488** | **400-590** | **7** | **600** | **72** |
| $P_{43}\text{-}mKate2$ | **561** | **590-700** | **2.5** | **600** | **86** |
| **BM1625 and BM1626** |  |  |  |  |  |
| $P_{spoIIQ}\text{-}yfp$ | **488** | **400-590** | **7** | **600** | **72** |
| $P_{43}\text{-}mKate2$ | **561** | **590-700** | **2.5** | **600** | **86** |

Fluorescence images were taken with an AxioCam MRm Rev.3 camera. The captured images were analyzed using the Zen Blue 2.3 lite software (Zeiss, Göttingen, Germany). Pixels showing P_*43*_ constitutive expression were normalized per maximum fluorescence intensity of each Z-stack. All other pixel values showing transcription from other promoters (P_*epsA*_, P_*tapA*_, P_*spoIIQ*_) are raw values, without normalization per maximum value. Z-stacks are presented as orthogonal displays, showing the stack exhibiting the highest fluorescence intensity.

### 1.6.7 Statistical analysis

All experiments were performed in at least three time independent biological replicates. We subtracted the estimate for background fluorescence intensity (averaged from all the wells with strains without fluorescent transcriptional reporter fusions) from every microtiter plate well fluorescence measurements. Afterwards, each well fluorescence intensity was averaged and a 95 % confidence interval was calculated from all of the wells. When performing biochemical tests and measuring CFUs all three independent biological replicates are shown with a straight line denoting their averages. Two groups of samples were compared by calculating a Student’s t-test and a one way non-parametric Mann-Whitney U test. We treated the two groups of samples as being statistically different if both tests showed a P-value < 0.05.

## 1.7 ACKNOWLEDGMENTS

We would like to acknowledge the ARRS (Slovenian Research Agency) P4-0116 program grant, the J4-9302 research grant and a young researcher grant awarded to M.S. We would also like to thank prof. dr. Nicola Stanley-Wall and prof. dr. Ákos T. Kovács for their inputs, debates and constructive criticism of the manuscript. This work has been supported by the infrastructural center “Microscopy of biological samples”, located in Biotechnical faculty, University of Ljubljana.

---

**Figure S1:**
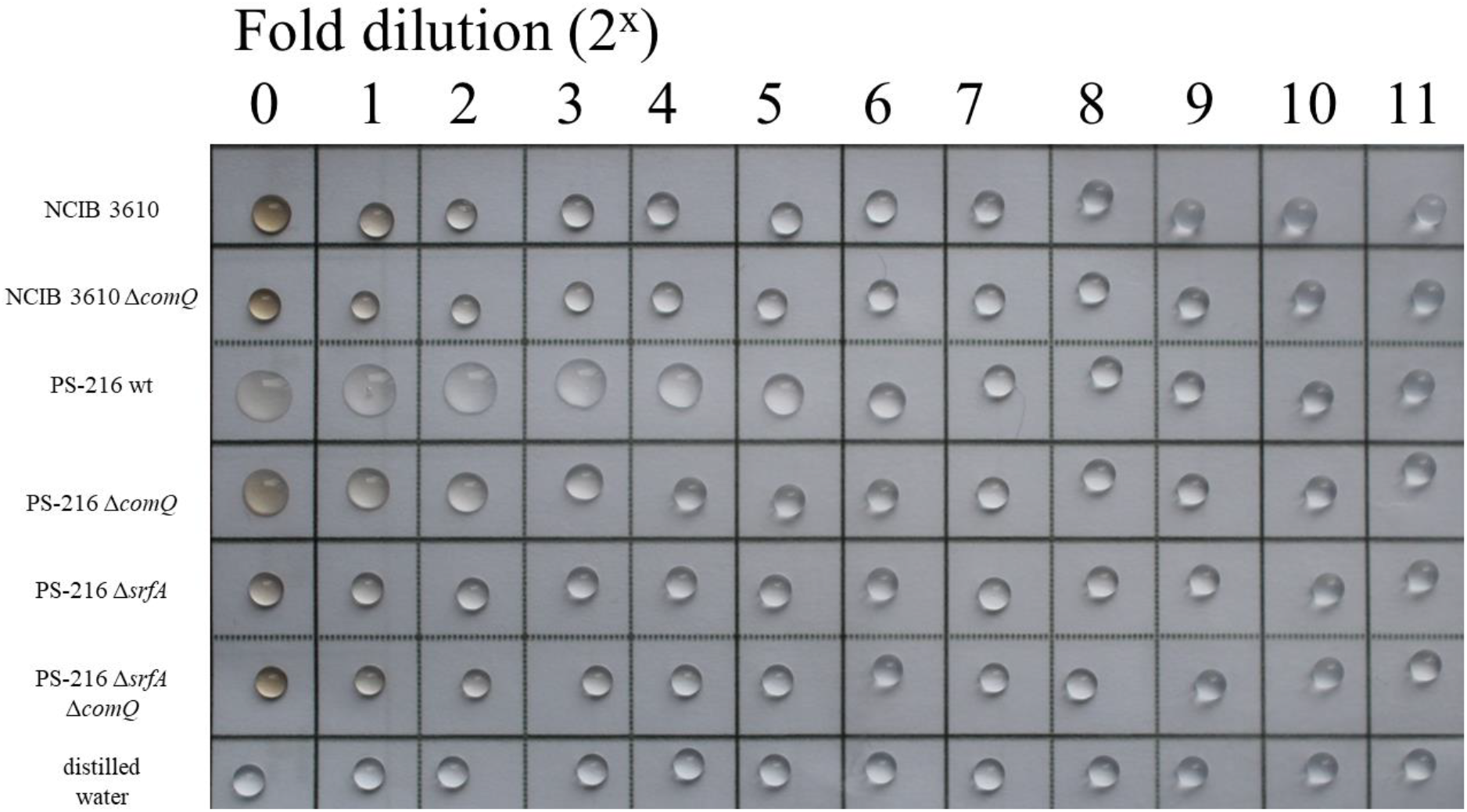
Semi quantification of surfactant concentrations in the floating biofilm spent media after 40 hours of static growth in MSgg medium at 37°C with the droplet surface wetting assay.

**Link to youtube video**

**Figure S2:** Fast forward video showing diffusion of the 50 μl methylene blue droplet placed on the PS-216 wild type (wt) or QS mutant (Δ*comQ*) floating biofilms grown statically in MSgg medium at 37 °C for the time periods indicated on the video.

**Figure S3:**
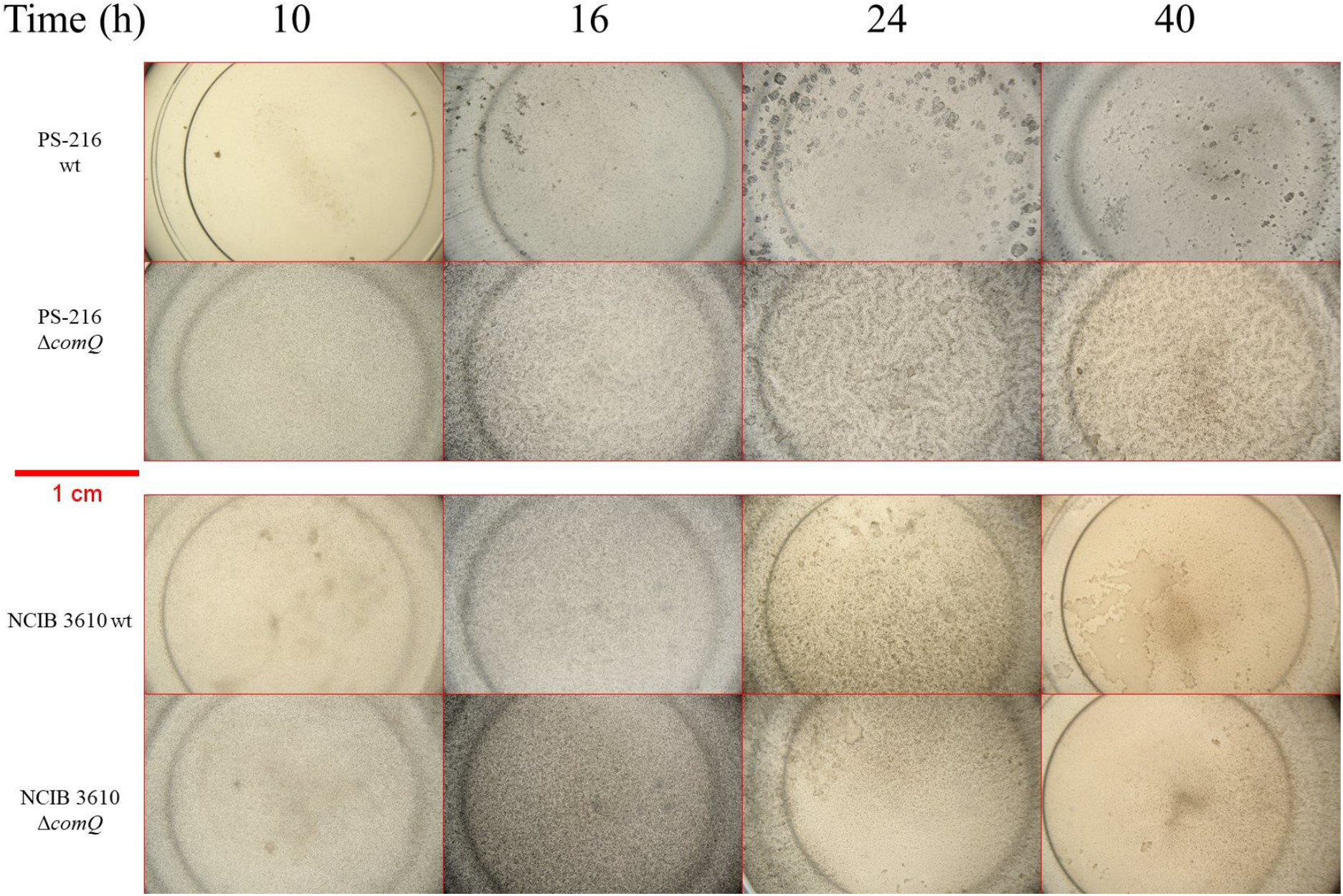
The formation of *B. subtilis* floating biofilms during static growth in MSgg medium at 37° C over time under 8-fold magnification.

**Figure S4:**
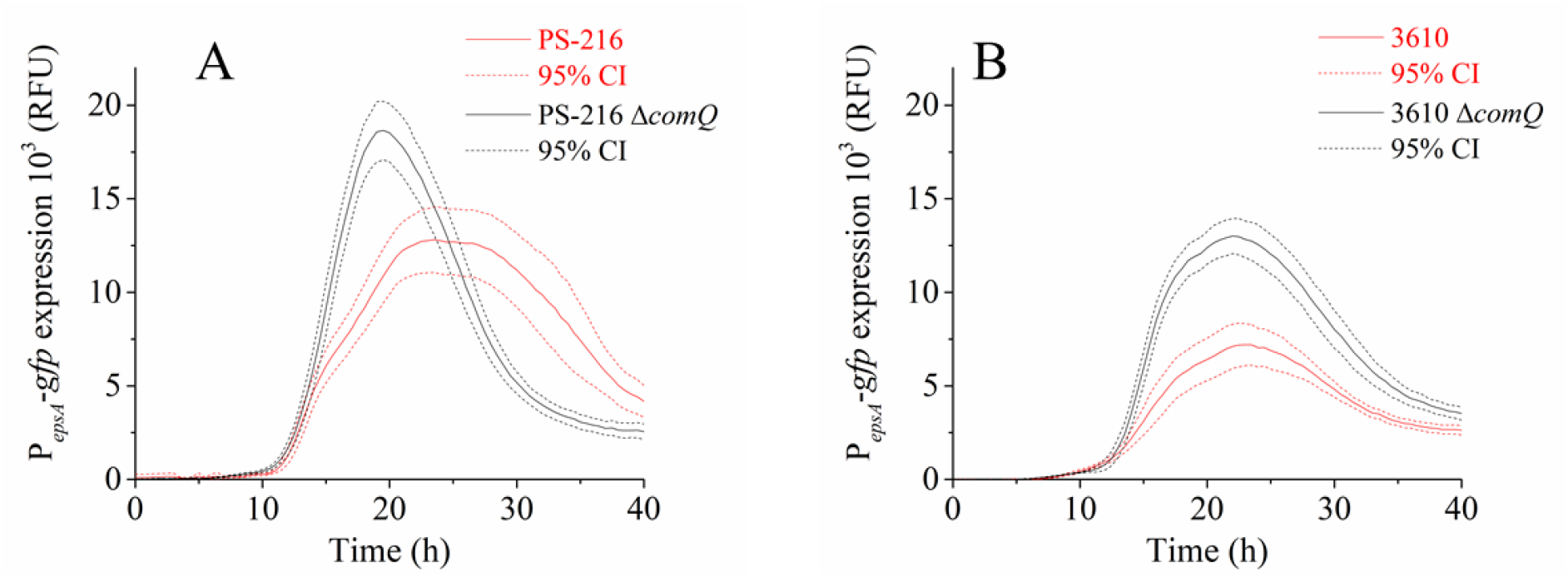
Cumulative P_*epsA*_-*gfp* promoter activity of *Bacillus subtilis* PS-216 wild type (wt) and QS mutant (Δ*comQ*) phenotypes (A) and in the NCIB 3610 wild type (wt) and QS mutant (Δ*comQ*) phenotypes (B) during static growth in MSgg medium at 37 °C.

**Figure S5:**
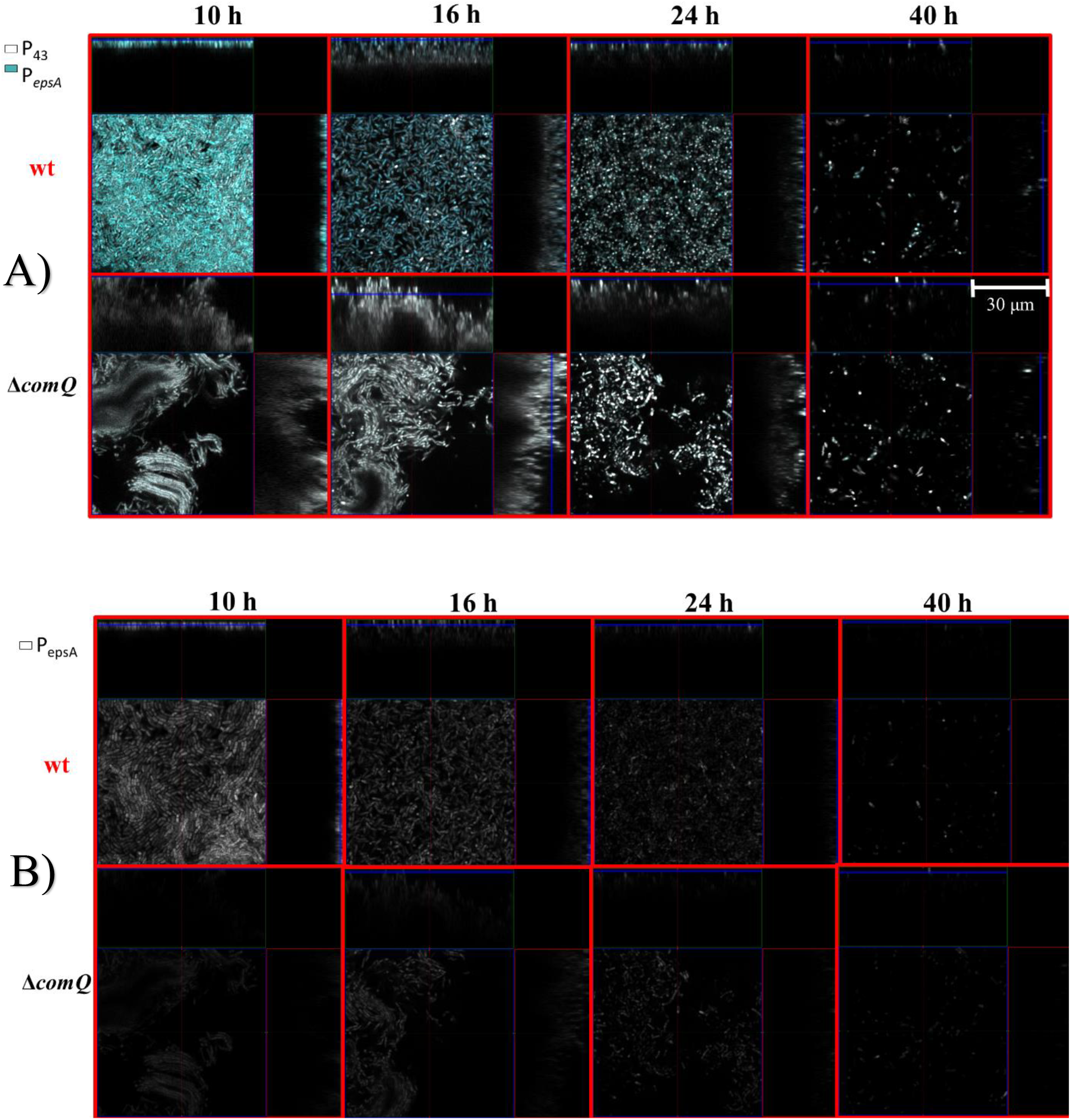
**A)** Confocal visualization of P_*43*_-*mKate2* fluorescence (false colored white; already shown in Figure 3) with the P_*epsA*_-*gfp* fluorescence (false colored teal) overlay in the wild type (wt) and QS mutant (Δ*comQ*) floating biofilms during static growth in MSgg medium at 37 °C. **B)** Same picture but showing only P_*epsA*_-gfp fluorescence.

**Figure S6:**
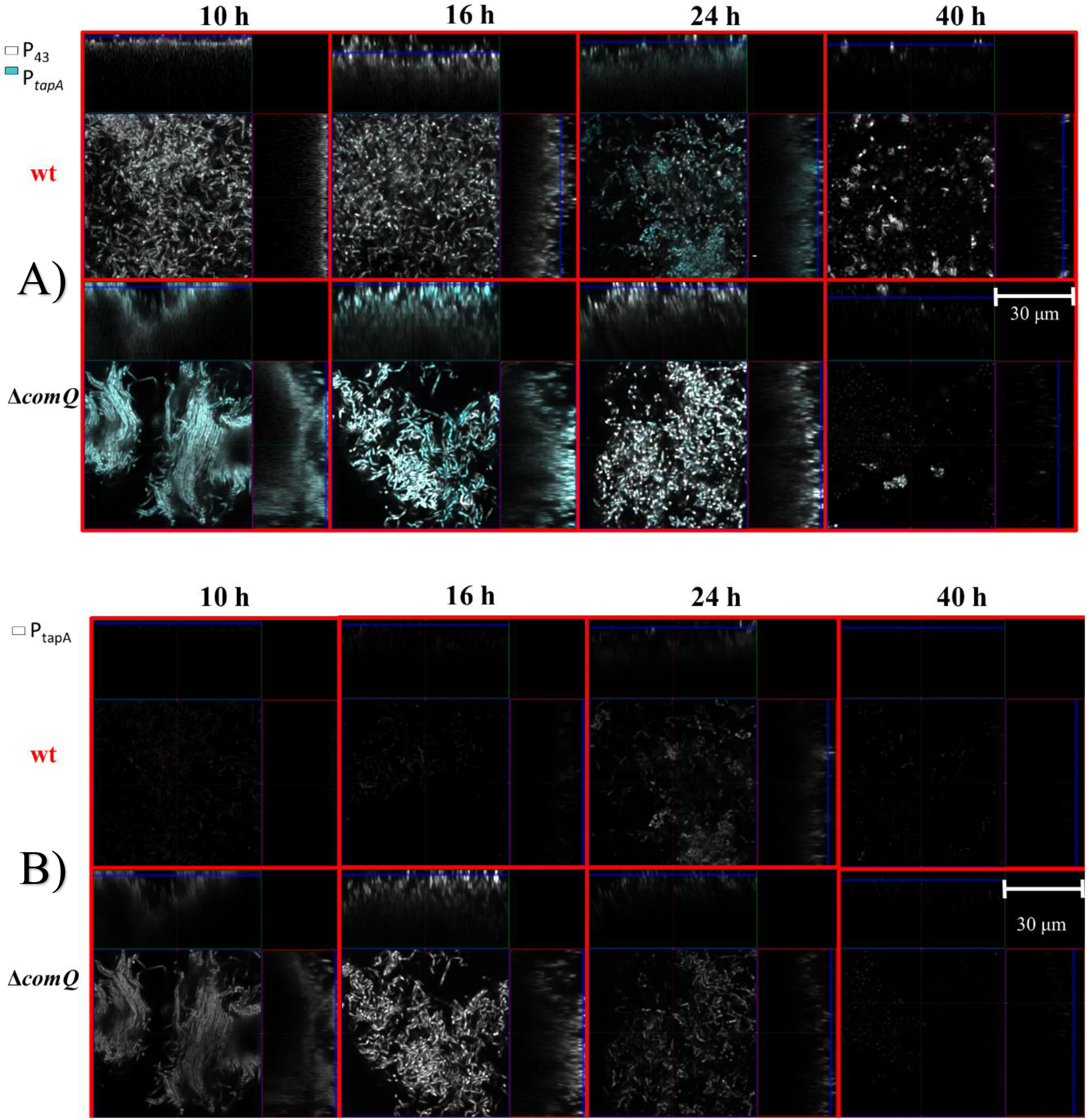
**A)** Confocal visualizations of P_*tapA*_-yfp fluorescence (false colored teal) and P_*43*_-mKate2 fluorescence (false colored white) expression in the wild type (wt) and QS mutant (Δ*comQ*) floating biofilms during static growth in MSgg medium at 37 °C. **B)** Same picture but showing only P_*tapA*_-yfp fluorescence.

## REFERENCES

1. Hall-Stoodley L, Costerton JW, Stoodley P. 2004. Bacterial biofilms: from the Natural environment to infectious diseases. Nat Rev Microbiol 2:95–108.

2. Jefferson K. 2004. What drives bacteria to produce a biofilm? FEMS Microbiol Lett 236:163–173.

3. Flemming H-C, Wingender J. 2010. The biofilm matrix. Nat Rev Microbiol 8:623–633.

4. Bloom-Ackermann Z, Ganin H, Kolodkin-Gal I. 2016. Quorum-sensing Cascades Governing Bacterial Multicellular Communities. Isr J Chem 56:302–309.

5. Platt TG, Fuqua C. 2010. What’s in a name? The semantics of quorum sensing. Trends Microbiol 18:383–387.

6. Mandic-mulec I, Stefanic P, Elsas JD V. 2015. Ecology of Bacillaceae. Microbiol Spectr 1–24.

7. Martinez RM. 2013. Bacillus subtilis. Brenner’s Encycl Genet Second Ed xxx:246–248.

8. Piggot PJ, Coote JG. 1976. Genetic aspects of bacterial endospore formation. Bacteriol Rev 40:908–62.

9. Kobayashi K. 2007. Gradual activation of the response regulator DegU controls serial expression of genes for flagellum formation and biofilm formation in Bacillus subtilis. Mol Microbiol 66:395–409.

10. Cairns LS, Hobley L, Stanley-Wall NR. 2014. Biofilm formation by *B acillus subtilis*: new insights into regulatory strategies and assembly mechanisms. Mol Microbiol 93:587–598.

11. Dragoš A, Kiesewalter H, Martin M, Hsu CY, Hartmann R, Wechsler T, Eriksen C, Brix S, Drescher K, Stanley-Wall N, Kümmerli R, Kovács ÁT. 2018. Division of Labor during Biofilm Matrix Production. Curr Biol 28:1903–1913.e5.

12. Abee T, Kovács ÁT, Kuipers OP, van der Veen S. 2011. Biofilm formation and dispersal in Gram-positive bacteria. Curr Opin Biotechnol 22:172–179.

13. López D, Kolter R. 2010. Extracellular signals that define distinct and coexisting cell fates in Bacillus subtilis. FEMS Microbiol Rev 34:134–149.

14. Veening J-W, Smits WK, Kuipers OP. 2008. Bistability, Epigenetics, and Bet-Hedging in Bacteria. Annu Rev Microbiol 62:193–210.

15. Schultz D, Wolynes PG, Jacob E Ben, Onuchic JN. 2009. Deciding fate in adverse times: Sporulation and competence in Bacillus subtilis. Proc Natl Acad Sci 106:21027–21034.

16. Lopez, D.; Vlamakis, H.; Kolter R. 2010. Biofilms Cold Spring Harbor Perspectives in Biology. Biofilms Cold Spring Harb Perspect Biol.

17. Kalamara M, Spacapan M, Mandic-Mulec I, Stanley-Wall NR. 2018. Social behaviours by Bacillus subtilis: quorum sensing, kin discrimination and beyond. Mol Microbiol 110:863–878.

18. Tran LSP, Nagai T, Itoh Y. 2000. Divergent structure of the ComQXPA quorum-sensing components: Molecular basis of strain-specific communication mechanism in Bacillus subtilis. Mol Microbiol.

19. Tortosa P, Logsdon L, Kraigher B, Itoh Y, Mandic-Mulec I, Dubnau D. 2001. Specificity and genetic polymorphism of the Bacillus competence quorum-sensing system. J Bacteriol 183:451–460.

20. Dogsa I, Choudhary KS, Marsetic Z, Hudaiberdiev S, Vera R, Pongor S, Mandic-Mulec I. 2014. ComQXPA quorum sensing systems may not be unique to Bacillus subtilis: A census in prokaryotic genomes. PLoS One 9.

21. Ansaldi M, Marolt D, Stebe T, Mandic-Mulec I, Dubnau D. 2002. Specific activation of the Bacillus quorum-sensing systems by isoprenylated pheromone variants. Mol Microbiol 44:1561–1573.

22. D’Souza C, Nakano MM, Zuber P. 1994. Identification of comS, a gene of the srfA operon that regulates the establishment of genetic competence in Bacillus subtilis. Proc Natl Acad Sci U S A 91:9397–9401.

23. Oslizlo A, Stefanic P, Dogsa I, Mandic-Mulec I. 2014. Private link between signal and response in Bacillus subtilis quorum sensing. Proc Natl Acad Sci 111:1586–1591.

24. Spacapan M, Danevčič T, Mandic-Mulec I. 2018. ComX-Induced Exoproteases Degrade ComX in Bacillus subtilis PS-216. Front Microbiol 9:1–23.

25. Msadek T, Kunst F, Klier A, Rapoport G. 1991. DegS-DegU and ComP-ComA modulator-effector pairs control expression of the Bacillus subtilis pleiotropic regulatory gene degQ. J Bacteriol 173:2366–2377.

26. Weinrauch Y, Msadek T, Kunst F, Dubnau D. 1991. Sequence and properties of comQ, a new competence regulatory gene of Bacillus subtilis. J Bacteriol 173:5685–5693.

27. Schneider KB, Palmer TM, Grossman AD. 2002. Characterization of comQ and comX, Two Genes Required for Production of ComX Pheromone in Bacillus subtilis. J Bacteriol 184:410–419.

28. Roggiani M, Dubnau D. 1993. ComA, a phosphorylated response regulator protein of Bacillus subtilis, binds to the promoter region of srfA. J Bacteriol 175:3182–3187.

29. Ogura M, Yamaguchi H, Ki Y, Fujita Y, Tanaka T. 2001. DNA microarray analysis of Bacillus subtilis DegU, ComA and PhoP regulons: an approach to comprehensive analysis of B.subtilis two-component regulatory systems. Nucleic Acids Res 29:3804–3813.

30. Comella N, Grossman AD. 2005. Conservation of genes and processes controlled by the quorum response in bacteria: characterization of genes controlled by the quorum-sensing transcription factor ComA in Bacillus subtilis. Mol Microbiol 57:1159–74.

31. Wolf D, Rippa V, Mobarec JC, Sauer P, Adlung L, Kolb P, Bischofs IB. 2015. The quorum-sensing regulator ComA from Bacillus subtilis activates transcription using topologically distinct DNA motifs. Nucleic Acids Res 1–13.

32. Ishii H, Tanaka T, Ogura M. 2013. The Bacillus subtilis Response Regulator Gene degU Is Positively Regulated by CcpA and by Catabolite-Repressed Synthesis of ClpC 195:193–201.

33. Verhamme DT, Murray EJ, Stanley-Wall NR. 2009. DegU and Spo0A jointly control transcription of two loci required for complex colony development by Bacillus subtilis. J Bacteriol.

34. Dogsa I, Brloznik M, Stopar D, Mandic-Mulec I. 2013. Exopolymer Diversity and the Role of Levan in Bacillus subtilis Biofilms. PLoS One 8:2–11.

35. Branda SS, González-Pastor JE, Ben-Yehuda S, Losick R, Kolter R. 2001. Fruiting body formation by Bacillus subtilis. Proc Natl Acad Sci U S A 98:11621–6.

36. Kearns DB, Chu F, Branda SS, Kolter R, Losick R. 2005. A master regulator for biofilm formation by Bacillus subtilis. Mol Microbiol 55:739–749.

37. Terra R, Stanley-Wall NR, Cao G, Lazazzera BA. 2012. Identification of Bacillus subtilis SipW as a Bifunctional Signal Peptidase That Controls Surface-Adhered Biofilm Formation. J Bacteriol 194:2781–2790.

38. Hamon MA, Stanley NR, Britton RA, Grossman AD, Lazazzera BA. 2004. Identification of AbrB-regulated genes involved in biofilm formation by Bacillus subtilis. Mol Microbiol 52:847–860.

39. Vlamakis H, Chai Y, Beauregard P, Losick R, Kolter R. 2013. Sticking together: building a biofilm the Bacillus subtilis way. Nat Rev Microbiol 11:157–168.

40. Jiang M, Shao W, Perego M, Hoch JA. 2000. Multiple histidine kinases regulate entry into stationary phase and sporulation in Bacillus subtilis. Mol Microbiol 38:535–542.

41. Chai Y, Chu F, Kolter R, Losick R. 2008. Bistability and biofilm formation in Bacillus subtilis. Mol Microbiol 67:254–263.

42. Shafikhani SH, Mandic-Mulec I, Strauch MA, Smith I, Leighton T. 2002. Postexponential Regulation of sin Operon Expression in Bacillus subtilis. J Bacteriol 184:564–571.

43. Bai U, Mandic-Mulec I, Smith I. 1993. SinI modulates the activity of SinR, a developmental switch protein of Bacillus subtilis, by protein-protein interaction. Genes Dev 7:139–148.

44. Chu F, Kearns DB, Branda SS, Kolter R, Losick R. 2006. Targets of the master regulator of biofilm formation in Bacillus subtilis. Mol Microbiol 59:1216–1228.

45. Kobayashi K, Iwano M. 2012. BslA(YuaB) forms a hydrophobic layer on the surface of Bacillus subtilis biofilms. Mol Microbiol 85:51–66.

46. Verhamme DT, Kiley TB, Stanley-Wall NR. 2007. DegU co-ordinates multicellular behaviour exhibited by Bacillus subtilis. Mol Microbiol 65:554–568.

47. Stanley NR, Lazazzera BA. 2005. Defining the genetic differences between wild and domestic strains of Bacillus subtilis that affect poly-γ-dl-glutamic acid production and biofilm formation. Mol Microbiol 57:1143–1158.

48. Marlow VL, Porter M, Hobley L, Kiley TB, Swedlow JR, Davidson FA, Stanley-Wall NR. 2014. Phosphorylated DegU manipulates cell fate differentiation in the Bacillus subtilis biofilm. J Bacteriol 196:16–27.

49. Lopez D, Fischbach MA, Chu F, Losick R, Kolter R, Lo D, Lopez D, Fischbach MA, Chu F, Losick R, Kolter R, Lo D, Lopez D, Fischbach MA, Chu F, Losick R, Kolter R. 2009. Structurally diverse natural products that cause potassium leakage trigger multicellularity in Bacillus subtilis. Proc Natl Acad Sci 106:280–285.

50. López D, Vlamakis H, Losick R, Kolter R. 2009. Paracrine signaling in a bacterium. Genes Dev 23:1631–1638.

51. Gallegos-Monterrosa R, Mhatre E, Kovács Ákos T. 2016. Specific Bacillus subtilis 168 variants form biofilms on nutrient-rich medium. Microbiol (United Kingdom) 162:1922–1932.

52. Shank EA, Kolter R. 2011. Extracellular signaling and multicellularity in Bacillus subtilis. Curr Opin Microbiol 14:741–747.

53. Dervaux J, Magniez JC, Libchaber A. 2014. On growth and form of Bacillus subtilis biofilms. Interface Focus 4:20130051–20130051.

54. Song Y, Nikoloff JM, Fu G, Chen J, Li Q, Xie N, Zheng P, Sun J, Zhang D. 2016. Promoter Screening from Bacillus subtilis in Various Conditions Hunting for Synthetic Biology and Industrial Applications. PLoS One 11:e0158447.

55. Doan T, Marquis KA, Rudner DZ. 2005. Subcellular localization of a sporulation membrane protein is achieved through a network of interactions along and across the septum 55:1767–1781.

56. Magnuson R, Solomon J, Grossman AD. 1994. Biochemical and genetic characterization of a competence pheromone from B. subtilis. Cell 77:207–16.

57. Nakano MM, Xia L, Zuber P. 1991. Transcription initiation region of the srfA operon, which is controlled by the comP-comA signal transduction system in Bacillus subtilis. J Bacteriol 173:5487–5493.

58. van Gestel J, Vlamakis H, Kolter R. 2015. New Tools for Comparing Microscopy Images: Quantitative Analysis of Cell Types in Bacillus subtilis. J Bacteriol 197:699–709.

59. Durrett R, Miras M, Mirouze N, Narechania A, Mandic-Mulec I, Dubnau D. 2013. Genome Sequence of the Bacillus subtilis Biofilm-Forming Transformable Strain PS216. Genome Announc 1.

60. Schlichting CD, Smith H. 2002. Phenotypic plasticity: linking molecular mechanisms with evolutionary outcomes 189–211.

61. Chastanet A, Vitkup D, Yuan G-C, Norman TM, Liu JS, Losick RM. 2010. Broadly heterogeneous activation of the master regulator for sporulation in Bacillus subtilis. Proc Natl Acad Sci 107:8486–8491.

62. Lowery NV, McNally L, Ratcliff WC, Brown SP. 2017. Division of labor, bet hedging, and the evolution of mixed biofilm investment strategies. MBio 8.

63. Veening J-W, Stewart EJ, Berngruber TW, Taddei F, Kuipers OP, Hamoen LW. 2008. Bet-hedging and epigenetic inheritance in bacterial cell development. Proc Natl Acad Sci 105:4393–4398.

64. Auchtung JM, Lee C a., Grossman AD. 2006. Modulation of the ComA-dependent quorum response in Bacillus subtilis by multiple rap proteins and Phr peptides. J Bacteriol 188:5273–5285.

65. Lazazzera BA, Solomon JM, Grossman AD. 1997. An Exported Peptide Functions Intracellularly to Contribute to Cell Density Signaling in B. subtilis. Cell 89:917–925.

66. Bischofs IB, Hug JA, Liu AW, Wolf DM, Arkin AP. 2009. Complexity in bacterial cell-cell communication: Quorum signal integration and subpopulation signaling in the Bacillus subtilis phosphorelay. Proc Natl Acad Sci 106:6459–6464.

67. Asfahl KL, Schuster M. 2017. Social interactions in bacterial cell–cell signaling. FEMS Microbiol Rev 41:92–107.

68. Van Gestel J, Weissing FJ, Kuipers OP, Kovács KT. 2069. Density of founder cells affects spatial pattern formation and cooperation in Bacillus subtilis biofilms. ISME J 852:2069–2079.

69. Otto SB, Martin M, Schäfer D, Hartmann R, Drescher K, Brix S, Dragoš A, Kovács ÁT. 2019. Privatization of biofilm matrix in structurally heterogeneous biofilms. bioRxiv 742593.

70. Lyons NA, Kolter R. 2017. Bacillus subtilis Protects Public Goods by Extending Kin Discrimination to Closely Related Species. MBio 8.

71. West SA, Diggle SP, Buckling A, Gardner A, Griffin AS. 2007. The Social Lives of Microbes. Annu Rev Ecol Evol Syst 38:53–77.

72. Pollak S, Omer-Bendori S, Even-Tov E, Lipsman V, Bareia T, Ben-Zion I, Eldar A, Bareia T, Omer-Bendori S, Lipsman V, Even-Tov E, Eldar A. 2016. Facultative cheating supports the coexistence of diverse quorum-sensing alleles. Proc Natl Acad Sci 113:2152–2157.

73. Albano M, Hahn J, Dubnau D. 1987. Expression of competence genes in Bacillus subtilis. J Bacteriol 169:3110–3117.

74. Oslizlo A, Stefanic P, Vatovec S, Beigot Glaser S, Rupnik M, Mandic-Mulec I. 2015. Exploring ComQXPA quorum-sensing diversity and biocontrol potential of Bacillus spp. isolates from tomato rhizoplane. Microb Biotechnol 8:527–540.

75. Stefanic P, Kraigher B, Lyons NA, Kolter R, Mandic-Mulec I. 2015. Kin discrimination between sympatric Bacillus subtilis isolates. Proc Natl Acad Sci 112:14042–14047.

76. Patrick JE, Kearns DB. 2008. MinJ (YvjD) is a topological determinant of cell division in Bacillus subtilis. Mol Microbiol 70:1166–1179.

77. Middleton R, Hofmeister A. 2004. New shuttle vectors for ectopic insertion of genes into Bacillus subtilis. Plasmid 51:238–245.

78. Norman TM, Lord ND, Paulsson J, Losick R. 2015. Stochastic Switching of Cell Fate in Microbes. Annu Rev Microbiol 69:381–403.

79. Stefanic P, Mandic-Mulec I. 2009. Social interactions and distribution of bacillus subtilis pherotypes at microscale. J Bacteriol 191:1756–1764.

80. Parashar V, Konkol MA, Kearns DB, Neiditch MB. 2013. A plasmid-encoded phosphatase regulates Bacillus subtilis biofilm architecture, sporulation, and genetic competence. J Bacteriol 195:2437–48.

81. Cuesta G, Suarez N, Bessio MI, Ferreira F, Massaldi H. 2003. Quantitative determination of pneumococcal capsular polysaccharide serotype 14 using a modification of phenol-sulfuric acid method. J Microbiol Methods 52:69–73.

82. Bradford MM. 1976. A rapid and sensitive method for the quantitation of microgram quantities of protein utilizing the principle of protein-dye binding. Anal Biochem 72:248–254.

